# Predator-prey interactions in a warming world: the critical role of cold tolerance

**DOI:** 10.1101/2022.01.17.476522

**Authors:** Xuezhen Ge, Cortland K. Griswold, Jonathan A. Newman

## Abstract

Thermal tolerance mismatch within predator-prey systems may have profound effects on species population abundances and geographical distributions. To examine the generalized responses of a predator-prey system to climate change, we construct a biologically detailed stage-structured population dynamic model of interactions between ladybird beetles and aphids. We explore the model’s dynamics across the entire feasible parameter space of mean temperature and seasonality. Within this space, we explore different scenarios of predator and prey thermal tolerance mismatch to gain insight into how these thermal sensitivities affect the interacting species’ responses to climatic change. Our results indicate a predator’s cold tolerance has a larger effect on prey abundance than its heat tolerance. Mismatches between the predator’s and prey’s thermal tolerances also affect the species’ response to climate change. We identify three common patterns of species abundance across the feasible parameter space that relate to the type of thermal tolerance mismatches. Our study highlights the importance of understanding the complex interplay between climate change and species interactions.

## 1. Introduction

The changing global climate is likely to alter species interactions (Blois et al., 2013; Alexander et al., 2015). Over the past few decades, progress has been made in understanding how individual species’ physiology, demography, and spatial distributions respond to climate change (Chen et al., 2011; Machekano et al., 2018). However, studies of the effects of climate change on species interactions are still under studied (Alexander et al., 2015; Gilman, 2017). The direct effect of climate on individual species can affect species interactions, which can likewise alter individual species’ performance under climate change (Blois et al., 2013; Boukal et al., 2019). It is thus crucial to understand the complex interplay between climate change and species interactions.

Predation is a fundamental biotic interaction (Bianchi et al., 2006; Glen and Dickman, 2014). Climate change may modify the characteristics of predator and prey individually, and this may alter each species’ phenology and potentially cause mismatches in the timing of life history events between the species. Such phenological mismatches may lead to complex out-comes and make difficult the task of predicting species’ population responses to climate change (Gilman, 2017; Schmitz and Barton, 2014; Damien and Tougeron, 2019). Boukal et al. (2019) provided a conceptual framework that links temperature effects on individual species to species interactions, as well as outlined how recent advances have revealed the importance of species interactions in maintaining ecosystem stability and resilience in the face of climatic change.

Predator and prey may differ in their thermal tolerance limits. For example, Buxton et al. (2020) found that two notonectid predators (*Anisops sardea* and *Enithares chinai*) and one copepod predator (*Lovenula falcifera*) had lower *CT_min_* and *CT_max_* (i.e., were more cold tolerant but less heat tolerant) than the three mosquito prey, *Aedes aegypti, Anopheles quadriannulatus* and *Culex pipiens*. Pintanel et al. (2021) quantified the thermal tolerance mismatch in a predator-prey system comprising dragonfly species and anuran species and found predators always had higher maximum thermal tolerances than their prey.

Much recent evidence suggests that the thermal tolerance limits of insect species have profound effects on species population abundance and geographical distributions as temperature is one of the most important abiotic constraints on species’ biological functions (Sunday et al., 2012; Birkett et al., 2018; Amundrud and Srivastava, 2020). Different thermal performances between predator and prey adds uncertainty and complexity when predicting species population abundance and distribution. Fig. 1 is a conceptual depiction of how thermal tolerance mismatches between interacting species result in different performance in response to climate change (e.g., wider thermal breadth of the predator may lead to a stronger predation rate). Despite increasing attention to the impacts of climate change on predator-prey interactions, few studies have evaluated how thermal tolerance mismatches between interacting species affect species’ response to a changing climate. Therefore, our goal in this work was to examine the generalized responses of a predator-prey system to climate change by incorporating different scenarios of predator and prey thermal tolerance mismatches (mainly focusing on horizontal shifts of thermal performance curves).

**Figure 1:**
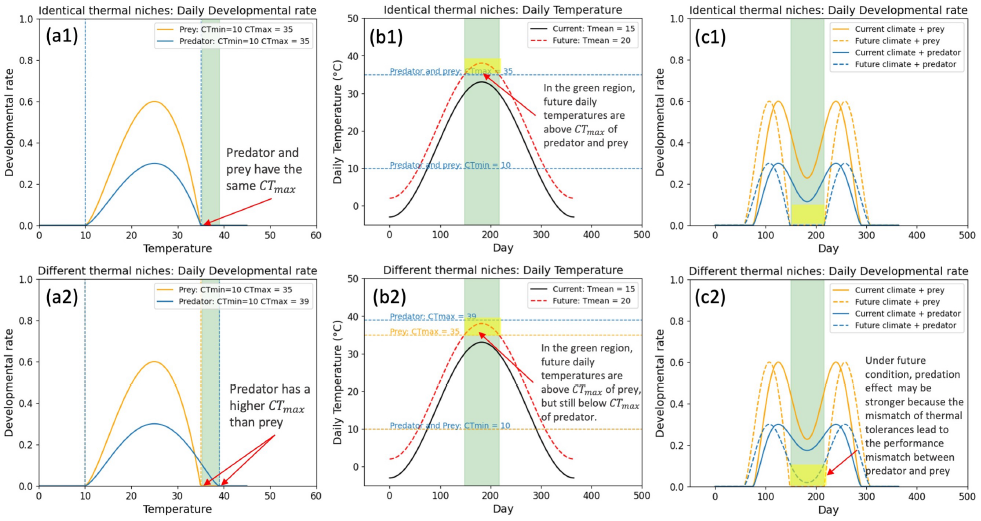
A demonstration of how thermal tolerance mismatches between interacting species result in different performances in response to climate change. (a1) and (a2) show the temperature performance curves for prey and predator’s developmental rates when they have identical (a1) or different (a2) thermal niches (i.e., predator is more heat tolerant than the prey). (b1) and (b2) show the interaction between the thermal tolerances and the changing temperatures under climate change (e.g., rising temperature may be suitable for the predator but not suitable for the prey when predator is more heat tolerant than the prey). (c1) and (c2) show the developmental rate curves for predator and prey under current and future climates, which demonstrate that the thermal tolerance mismatch between the interacting species leads to species’ different performances in response to climate change.

To achieve our goal, we construct a biologically detailed stage-structured population dynamic model of the interactions between ladybird beetles and aphids. Aphids and ladybirds occur worldwide, with aphids being amongst the most destructive insect pests to cultivated plants and an important vector of plant pathogens (Ng and Perry, 2004; Dedryver et al., 2010). The response of aphids to global climate change is therefore of broad concern. Ladybird beetles predate aphids and can be a natural regulator of aphid populations. They are also used in biological control practices (Brown, 2004). As ectotherms, aphids and ladybirds are both highly sensitive to ambient temperature. Due to their economic importance, a great deal of life history data are available for these species. Thus, these species constitute a good model system for addressing our research interests. We parameterized the model using experimental data on the responses of developmental, fecundity, mortality and predation rates to temperature. We based these parameter estimates on the cotton aphid (*Aphis gossypii* Glover [Hemiptera: Aphididae]) and ladybird beetle (*Harmonia dimidiata* (Fab.) [Coccinellidae: Coleoptera]), and constructed a mechanistic model of aphid-ladybird population dynamics (See Supplement 1 for the life history of *A. gossypii* and *H. dimidiata*). We analyzed this model for different predator-prey thermal tolerance mismatch scenarios and climate scenarios to gain insight into how such mismatches for interacting species affects their responses to climate change and how climate change affects the population abundance for interacting species. Although our model is based on aphids and ladybirds, it is informative for species with similar interactions, as well as our current state of knowledge of the response of aphids to global warming.

## 2. Material and Methods

In this study, we develop a stage-structured population dynamic model of aphids and ladybirds. Developmental, fecundity, mortality and predation rates for both aphids and ladybirds are functions of temperature (Zamani et al., 2006; Khan et al., 2015). We use the model to estimate annual aphid pressure (*AAP*, defined as the accumulation of the daily population abundance) and annual ladybird pressure (*ALP*) for different temperature profiles and different predators. Note that, for brevity, we will refer *AAP* and *ALP* as aphid and ladybird population abundances throughout the paper, but we recognize that they are abundances across the entire year.

### 2.1. Predator and prey thermal tolerance mismatch scenarios

We used nine hypothetical ladybirds and one aphid to form nine pairs of interacting species which have all the possible qualitative thermal tolerance mismatches. The nine hypothetical ladybirds have different critical thermal minima and maxima. The thermal tolerances of the nine pairs of species are summarized in Table 1, and illustrated in Fig. 2. AL1 (aphid - ladybird1) is a base scenario, where the aphid and ladybird beetle have identical thermal niches. For the next three pairs of species (AL2, AL3 and AL4), ladybird beetles are *more* thermotolerant than the aphids (ladybirds have greater thermal breadth and either better cold or heat tolerance, or both). For the next three pairs (AL5, AL6 and AL7), ladybird beetles are *less* thermotolerant than the aphids (ladybirds have less thermal breadth and poor cold or heat tolerance, or both). For AL8 and AL9, ladybirds have the same thermal breadth as the aphid, but their cold and heat tolerances are different.

**Table 1:**
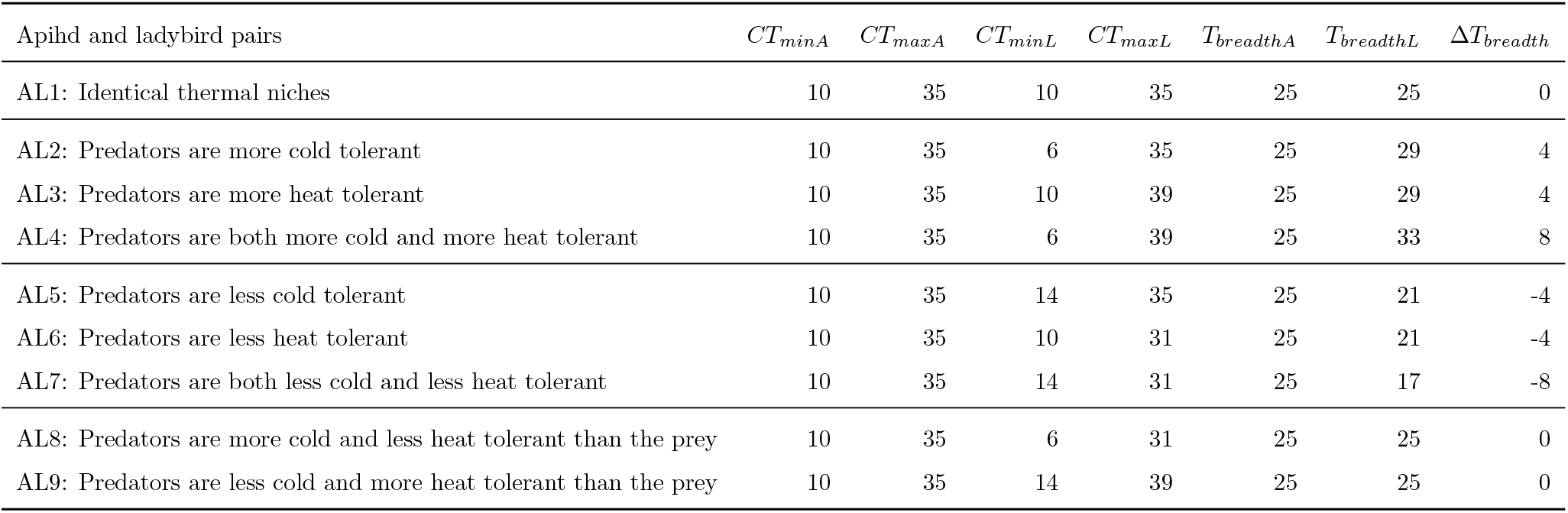
Thermal tolerances for the nine pairs of aphid and ladybird beetles. *CT_minA_*, *CT_maxA_*, *CT_minL_* and *CT_maxL_* represent critical thermal minimums and maximums for aphid and ladybirds. *T_breadthA_* = *CT_maxA_* – *CT_minA_*, *T_breadthL_* = *CT_maxL_* − *CT_minL_*, Δ*T_breadth_* = *T_breadthL_* − *T_breadthA_*.

**Figure 2:**
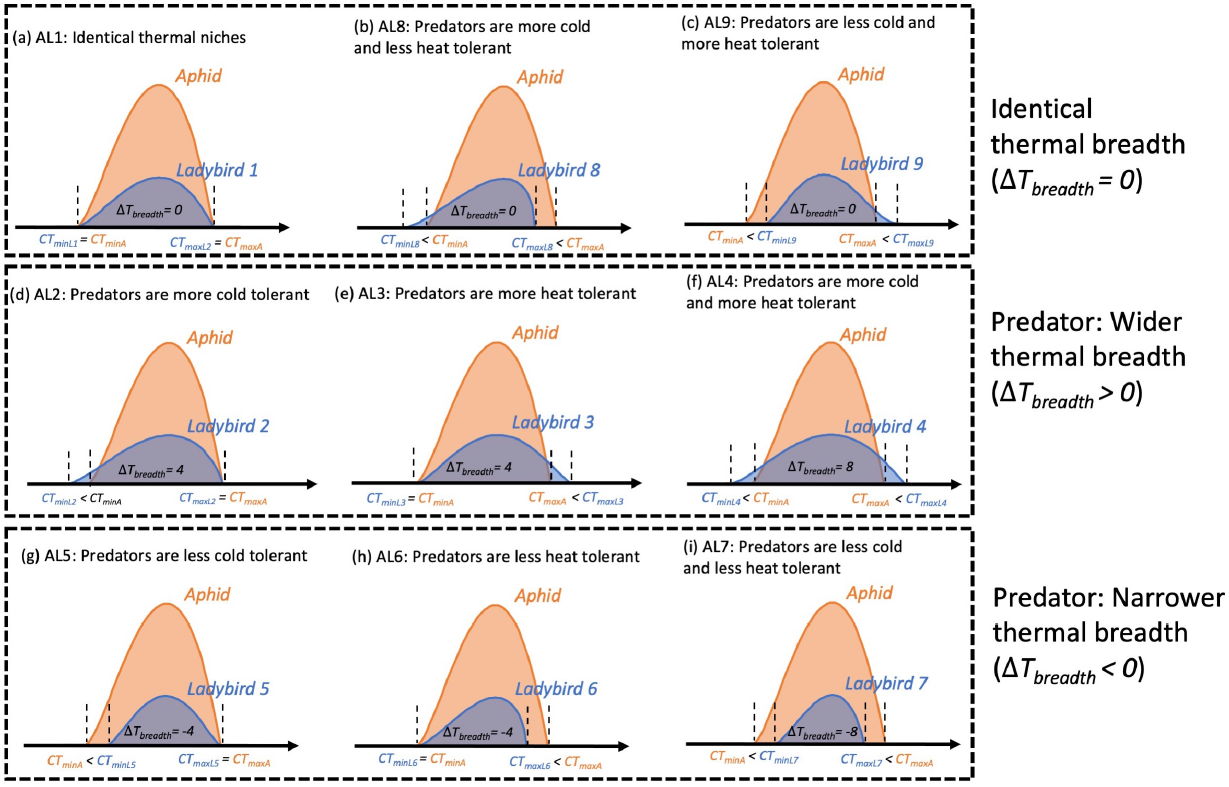
Conceptual sketch of thermal tolerance mismatches for nine pairs of aphid and ladybird beetles. The orange curve represents the thermal performance curve for the aphid, the blue curve represents the thermal performance curve for the ladybird. *CT_minA_*, *CT_maxA_*, *CT_minL_* and *CT_maxL_* represent critical thermal minima and maxima for aphid and ladybirds. Δ*T_breadth_* = *T_breadthL_* – *T_breadthA_*. A lower maximum thermal performance of ladybirds relative to aphids corresponds to a lower intrinsic developmental rate.

### 2.2. Model construction

Following the traditions of insect predator-prey modelling (see e.g. Xia et al. (2018)), we constructed a continuous time stage-structured model for aphid and ladybird population dynamics. We model developmental, birth, death, and predation rates as temperature dependent. The dynamic model comprises a set of coupled ordinary differential equations. The Python code for the model is available at https://github.com/xuezhenge/population-dynamic-model.

#### 2.2.1. Assumptions and Modelling Choices

1. We model only the anholocyclic life cycle and asexual reproduction of aphid.
2. For each locality, we assume that the aphid and ladybird populations are closed (e.g., no immigration and emmigration).
3. We assume the stage-specific temperature-dependent predation rates of ladybird are the same for all ladybird life stages.
4. For the aphid, we assume that predation by ladybirds is the only source of aphid extrinsic mortality. For the ladybird, we assume that ladybird birth rates depend on aphid capture rates, and that the only source of extrinsic mortality in our model derives from starvation at low aphid population sizes. Both aphid and ladybird intrinsic mortality rates are functions of temperature.
5. We assume *CT_minA_*, *CT_maxA_*, *CT_minL_* and *CT_maxL_* are the critical thermal minima and maxima for aphid and ladybird beetle, beyond which developmental and predation rates are set to 0, the mortality rates are maxima.
6. Although our model is stage-specific, we assume all the stages share the same temperature-dependent developmental rate and intrinsic mortality rate to simplify the analysis.

#### 2.2.2. General temperature-dependent function

We often make use of the same function to model temperature-dependence, albeit with differing values of the shape parameters. It is therefore convenient to define the function generally. We used a function presented by Thornley and France (2007, p. 105, their eqn 4.50) to model fecundity and development as functions of temperature.

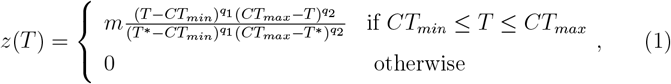

This function is flexible and fits the temperature-dependent traits very well. Eq. 1 has three shape parameters: *m* scales the function by changing its maximum value, and *q*_1_ and *q*_2_ change the function shape. *T* is the air temperature. The function reaches to its maximum (*m*) at *T*\*. *CT_min_* and *CT_max_* are the minimum and maximum temperature thresholds beyond which fecundity or developmental rate is 0, their values for different species pairs are provided in Table 1.

Eq. 1 reaches its maximum value at

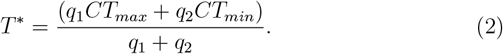

To focus on the effect of thermal tolerance limits and isolate the effect of altering thermal performance curve, we fix the value of *T*\* to be 25 °C, which is approximately the optimal temperature for the development of the aphid and ladybird. To ensure *T*\* stays constant for all species, we fix the value of *q*_1_ and solve Eq. 2 for *q*_2_.

We used a piecewise linear function, *υ*(*T*), to fit the experimental intrinsic mortality data. This function is flexible and has the correct general shape for describing how mortality rates change with temperature:

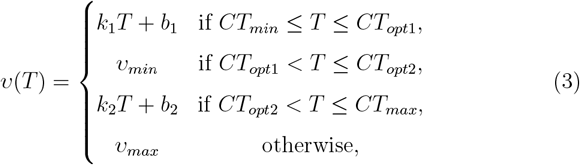

Here, *T* is the air temperature, *CT*_*opt*1_ and *CT*_*opt*2_ define the temperature range in which mortality is minimal, which we set to be 20 °C and 30 °C for all species. *CT_min_* and *CT_max_* are the minimum and maximum temperature thresholds beyond which mortality rates are maximal (see Table 1). *k*_1_, *b*_1_, *k*_2_ and *b*_2_ are the parameters which make *υ*(*CT*_*opt*1_) = *υ*(*CT*_*opt*2_) = *υ_max_* and *υ*(*CT_min_*) = *υ*(*CT_max_*) = *υ_min_*. We could not find empirical estimates for *υ_max_* in the literature. A sensitivity analysis showed that different maximal mortality rate of aphid (*υ_maxA_*) did not strongly affect the population dynamics of aphid and ladybird (Fig. S5), so we somewhat arbitrarily assumed *υ_maxA_* = 0.3, which is higher than the maximum mortality rates that were obtained from experiments. Similarly, we had to guess *υ_maxL_* due to insufficient experimental data. We picked *υ_maxL_* = 0.15 as ladybirds generally have a lower mortality rate than aphids. It is not surprising to see larger ladybird’s maximal mortality rate (*υ_maxL_*) drives larger aphid population due to fewer predators (Fig. S6). These values *υ_minA_* and *υ_minL_* for and seem compatible with the available experimental data (Van Steenis and El-Khawass, 1995; Zamani et al.,2006; Yu et al., 2013; Mou et al.,2015; Khan et al.,2015, 2016a,b).

#### 2.2.3. Aphid Submodel

Lacking good estimates of aphid overwinter survival, we model only the anholocyclic life cycle and asexual reproduction. We only keep track of the apterous adults in the aphid population, and assume that all the nymphs will develop to be apterous adults. Based on the life stages of *A. gossypii*, we model five stages, four instar nymphs and apterous adults. A schematic diagram of the aphid submodel (Fig. 3a) illustrates the dynamic processes of the model. There are four rate variables: fecundity, development, predation, and mortality. We know from laboratory experiments that these rates are all temperature-dependent (Aldyhim and Khalil, 1993; Van Steenis and El-Khawass, 1995; Kersting et al.,1999; Xia et al., 1999; Satar et al., 2005; Zamani et al., 2006; Singh and Singh, 2015). We therefore model these rates as functions of air temperature. The mathematical notations used in this submodel are summarized in Table S1. All the estimated parameter values for the aphid are listed in Table S2, which we derived from *A. gossypii* life history data to get the correct general shape for temperature performance curves. The shapes of the temperature-dependent fecundity rate, development rates and mortality rates of different stages of the nine pairs of species are shown in Fig. S1.

**Figure 3:**
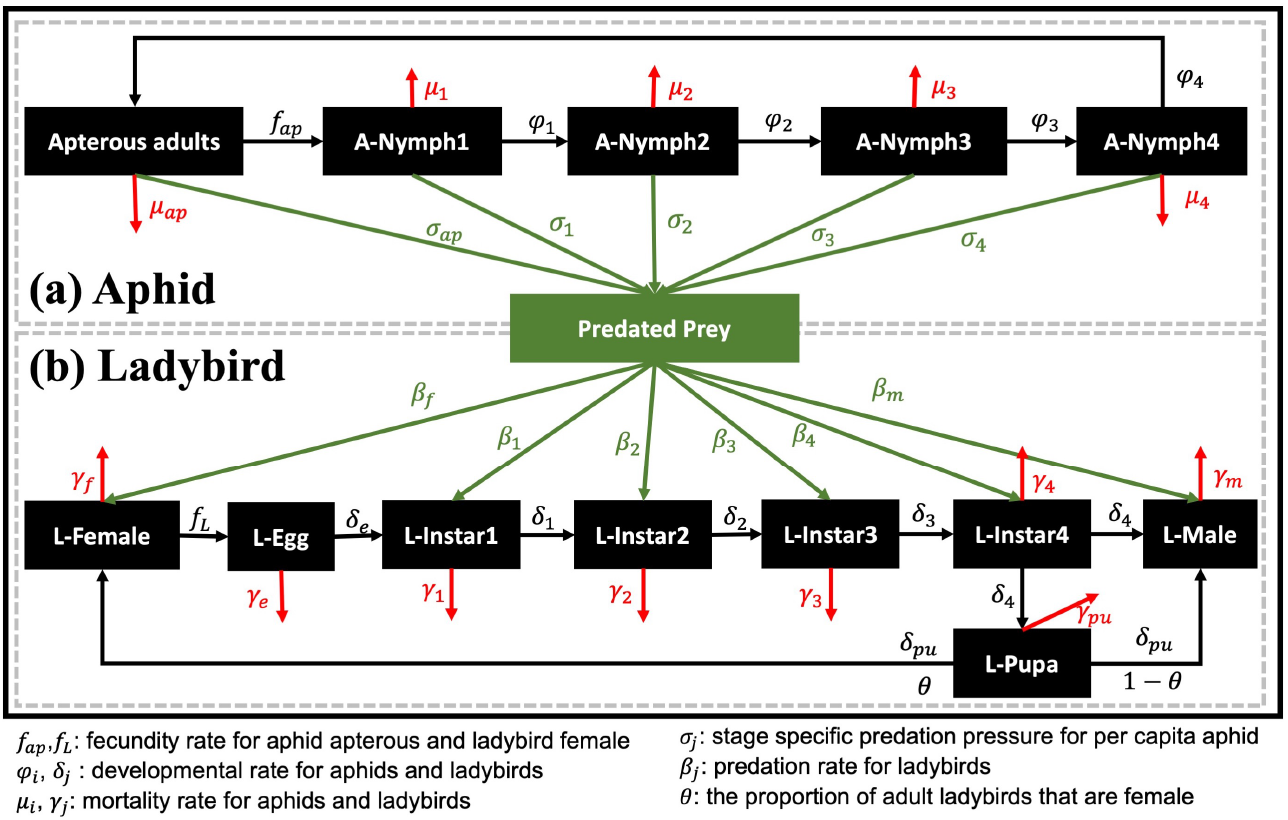
Schematic diagrams of model structure and analysis. (a) and (b) denote the life histories of the aphid and ladybird beetle, respectively. They are connected by predation. The black boxes in (a) and (b) represent different life stages of the two species. The parameters represent development rates, fecundity rates, mortality rates and predation rates (see Table S1 and S3 for the definitions of these parameters.)

##### Fecundity rate of apterous aphid adults

Since we only consider asexual reproduction of apterous adults in our model, we use the fecundity rate (*f_ap_*) to denote the total number of nymphs produced per apterous adult per day. The per capita fecundity rate of apterous adults is given by Eq. 1 where *f_ap_*(*T*) ≡ *z*(*T*). At peak densities for *A. gossypii,* the highest density of aphids per leaf is around 200 (Slosser et al., 2004). We assume a planting density of 20,000 to 70,000 cotton plants per acre (= 4.94 to 17.30 plants m^−2^) and 30 leaves per plant. From this, we estimated the aphid carrying capacity (*K*) to be approximately 5 × 10^4^ aphids m^−2^.

##### Developmental rate of nymphs

The four instar stages have similar development times, thus for simplicity we use the same developmental rate for all instar nymphs (*φ*). The per capita developmental rate of the nymphs is give by Eq. 1 where *φ* (*T*) ≡ *z*(*T*) (see Table S2 for parameter values).

##### Mortality rate of apterous adults and nymphs

In the field, aphids experience both intrinsic and extrinsic mortality. Intrinsic mortality is assumed to be the result of biological aging, which can be estimated with the experimental data obtained in the lab with *ad libitum* food. However, extrinsic mortality, which results from environmental hazards and natural enemies, is exceedingly difficult to estimate experimentally. We assumed that the only source of extrinsic mortality in our model derives from ladybird beetle predation. The mortality rates of apterous adults and nymphs that we used in our model represent the per capita daily intrinsic mortality. The per capita mortality rate of apterous adults is given by Eq. 3 where *μ_ap_* (*T*) ≡ *υ*(*T*). For simplicity, we assumed nymphs and apterous adults have the same intrinsic mortality rates, i.e., *μ_ny_*(*T*) ≡ *μ_ap_*(*T*) (see Table S2 for parameter values).

##### Predation rates on aphid life stages

We found no empirical evidence suggesting that the ladybird has a preference for a specific aphid developmental stage so we assume that the ladybird beetle predates each stage of the aphid in proportion to its relative abundance in the total aphid population. Let *A_i_* be the stage-specific aphid local population size, and let *A_den_* be the total aphid population size, given by,

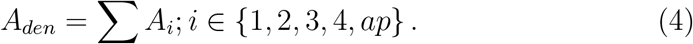

Then, let *α_i_* be the fraction of each stage in the total aphid population, given by,

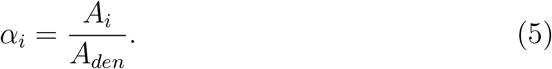

Finally, let *P* be the total number of aphids captured by the ladybird population per day (see Eq. 14). Therefore the number of stage specific predated aphids (*P_i_*) and the stage specific predation pressure for per capita aphid (*σ_i_*, i.e., probability of each aphid being predated by ladybirds) are given by,

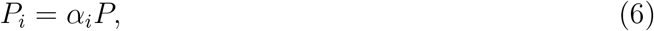

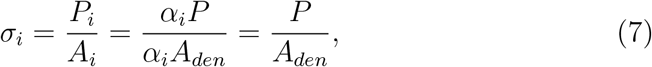

again, where *i* =1, …, 4, *ap* denote the four aphid nymphal instars and the apterous adults. *P_i_* represents the number of stage-specific aphids captured by the ladybid population per day and is a function of the per capita prey capture rate and the total population size of the ladybird beetles, as explained in the ladybird submodel section below (section 2.2.4, Eq. 14). Eq. 7 indicates that the predation pressure from ladybirds (*σ_i_*) is the same for each life stage of aphids based on our assumption that the ladybird beetle predates each stage of the aphid in accordance with its relative abundance in the total aphid population.

##### Aphid submodel state equations

The rates of change of the population of the various stages of aphid’s life history are modelled by the following coupled ordinary differential equations:

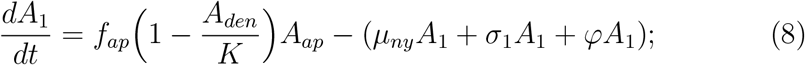

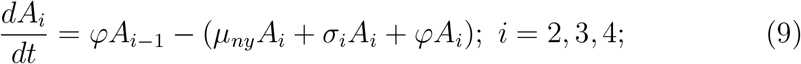

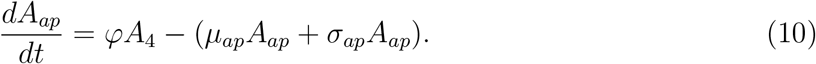

#### 2.2.4. Ladybird beetle submodel

There are eight state variables in the submodel of the ladybird, which corresponds to its life stage, including egg, four instar larvae, pupa, female and male adults. The dynamic process of the submodel is shown schematically in Fig. 3b. We consider four rate variables: fecundity, development, mortality, and predation. Since we assume that the ladybird beetle feeds only on the aphids, the number of captured aphids per ladybird per day (per capita predation rate) affects the ladybird’s fecundity and developmental rates. The mathematical notation used throughout the remainder of this submodel is shown in Table S3 and all the estimated parameter values for the nine ladybird beetles are listed in Table S4. All of these rate variables depend on temperature. Laboratory experiments of *Harmonia dimidiata* under various temperature conditions provided the life tables we used for estimating the parameters for the rate variables (Khan et al., 2015, 2016a,b; Sharma et al., 2017).

##### Predation rate

Except for the egg and pupa, all the other stages of the ladybird can predate aphids. The predation rate for different stages denotes the number of aphids eaten by per ladybird, per day. We assume that the ladybird’s predation rate depends on both aphid density and ambient air temperature.

###### *Predation rate as a function of aphid density at* optimal *temperatures*

Previous empirical studies indicated that a Type II functional response is common in coccinellids for all life stages of the ladybird (Zarghami et al., 2016; Sharma et al.,2017). We used Holling’s (1959) Type II functional response to describe the relationship between the number of consumed prey and the prey density. Let *β*(*A_den_*, *T_opt_*) be the predation rate per ladybird beetle as a function of prey density (*A_den_*) at the optimal temperature (*T_opt_*), and 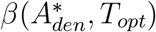 be the maximum intake rate at maximal aphid density 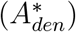, then,

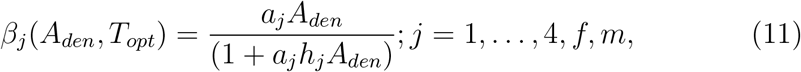

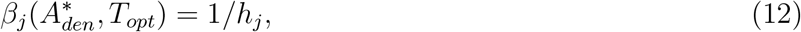

where in Eq. 11 *a_j_* and *h_j_* denote predator’s searching rate and handling time, respectively (Holling, 1959). Previous laboratory experiments have estimated the effect of aphid density on predation rates for different life stages of the ladybird (Agarwala et al., 2009; Sharma et al., 2017), thus the values of *a_j_* and *h_j_* were estimated from density-dependent predation data at the optimal temperature *T_opt_*.

###### *Predation rate as a function of temperature at* maximal *aphid density*

Laboratory experiments show that ladybird predation rates are also temperature-dependent (Yu et al.,2013; Khan et al., 2016a,b). Khan et al. (2016a) found that the relationship between predation rates of different stages of *H. dimidiata* and various temperatures are similar to each other, so we used a single temperature-dependent predation rate function for all life stages of ladybirds. Let 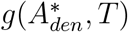 be the temperature-dependent predation rate at prey saturation 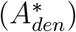, which is give by the general temperature dependence function (Eq. 1).

###### Predation rate as a function of both aphid density and temperature

Let *β*(*A_den_*, *T*) be the predation rate as a function of both density and temperature:

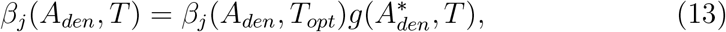

where *β*(*A_den_*, *T_opt_*) is given by Eq. 11.

The total rate of prey consumption by the predator population (*P*) can then be given by the following equation:

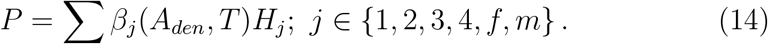

##### Fecundity rate of female ladybirds

The fecundity rate (*f_L_*) of female ladybirds represents the number of eggs produced per female per day, which depends on the females’ predation rate. The more prey consumed, the more energy the predator can allocate for reproduction. The predator’s numerical response (*Q_p_*; also called the ‘transformation rate’) is the mean number of aphids a ladybird needs to consume to reproduce a single egg. The fecundity rate is given by the following function:

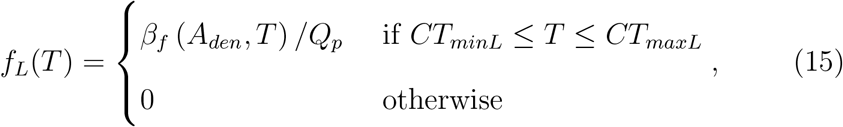

where *β_f_*(*A_den_*, *T*) is given by Eq. 13. We conducted a sensitivity analysis for *Q_p_*, which shows that annual aphid population and persistent times are sensitive to the choice of this parameter value when temperatures are warmer and relatively constant throughout the year (Fig. S3d1-d4). However, there are no sufficient data to estimate the temperature-dependent *Q_p_*. For simplicity, we assume *Q_p_* to be a consent value 100 aphids per ladybird, which is a rough guess based on Yu et al.’s (2013) laboratory experiment.

##### Stage-Specific Developmental rates

We assume that developmental rates depend on both the ambient air temperature and on the predation success of each specific ladybird life stage.

###### *Temperature-dependent development rates for* egg and pupa

The developmental rates of egg (*δ_∊_*) and pupa (*δ_p_*) are temperature-dependent and assumed to be the same, they are given by Eq. 1 where *δ_∊_*(*T*) ≡ *δ_p_*(*T*) ≡ *z*(*T*) (see Table S4 for parameter values).

##### *Temperature-dependent developmental rates for* larvae

###### Temperature-dependent developmental rates at prey saturation

The developmental rates of larvae (*δ_j_*) also depend on the stage-specific predation rates. The more prey consumed, the more energy the predator can use to support its development. We assume that the developmental rate for the various instars estimated from laboratory data represents the temperature-dependent developmental rate at prey saturation, 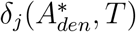, given by Eq. 1 where 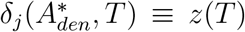 (*j* = 1, 2, 3,4) (see Table S4 for parameter values). The values of temperature-dependent developmental rates for larvae are assumed to be the same as for the egg and pupa.

###### Index that scales temperature-dependency and prey saturation

Recall, 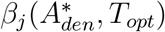 is the maximum prey intake rate (see Eq. 12) and *β_j_*(*A_den_*, *T*), given by Eq. 13, is the aphid-density and temperature-dependent predation rate. So, we use an index (*η*) that ranges from 0 to 1 and scales the temperature-dependent and prey saturated predation rate by the rate of prey-captured relative to the maximum prey-capture rate (1/*h_j_*), given by:

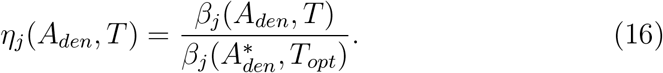

###### Development rates as a function of both prey saturation and temperature

Next, let *δ_j_*(*A_den_*, *T*) be the prey density-dependent and temperature-dependent development rate of larvae, given by:

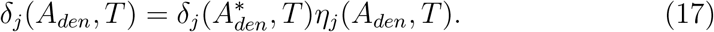

##### Mortality rate of various stages

We assume that the only source of extrinsic mortality in our model derives from starvation due to low aphid densities. The mortality rate of the ladybird (*γ_j_*) denotes the daily *intrinsic* mortality. The per capita mortality rate of different stages is given by Eq. 3 where *γ_j_* (*T*) ≡ *υ_i_*(*T*) (see Table S4 for parameter values), *j* = *∊*, 1, …4, *p*, *f*, *m* denote all the stages of the ladybird, and their mortality rates are assumed to be same.

##### Ladybird submodel state equations

The rates of change of the population of the various developmental stages of *H. dimidiata* were modelled by the following coupled ordinary differential equations:

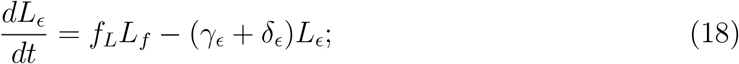

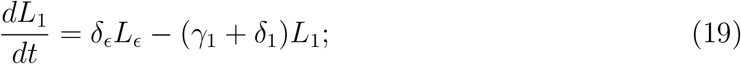

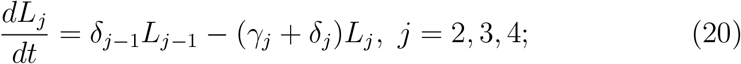

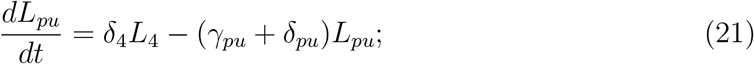

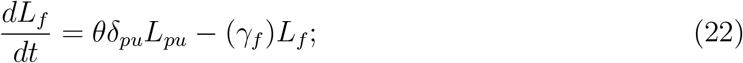

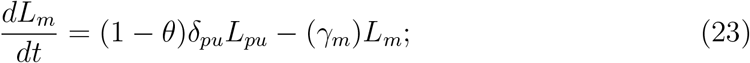

where *θ* represents the proportion of adult ladybirds that are females, which we assume is 0.5 (Farhadi et al., 2011).

### 2.3. Temperature profiles

For simplicity, we use a cosine function of daily mean temperature to generate various temperature profiles (Eq. 24). In Eq. 24, *s* and 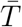 represent the amplitude and vertical shift respectively, and *t* denotes time (day). *s* represents the ‘seasonality’ of daily temperature, which is defined as half of the difference between minimum daily temperature (*T_yearmin_*) and maximum daily temperature (*T_yearmax_*) in a year (Eq. 25). 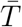 represents the yearly mean temperature (Eq. 26).

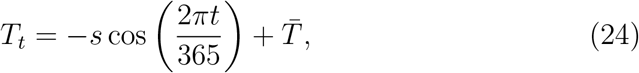

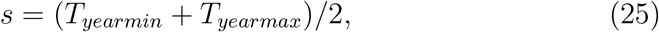

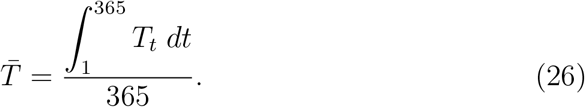

To obtain the feasible range of the two temperature metrics, seasonality (*s*) and yearly mean temperature 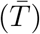, we calculate the local temperature metrics in 2020 and 2080 using the daily temperature data at 1° × 1° grid resolution. We downloaded the hourly 2-meter temperature in 2020 from ERA5 (https://www.ecmwf.int/en/forecasts/datasets/reanalysis-datasets/era5) and the daily temperature projections from the Coupled Model Intercomparison Project Phase 6 (CMIP6) (https://esgf-node.llnl.gov/search/cmip6/). EAR5 shows the hourly data of a global climate reanalysis from 1979 to the present (documentation is available at https://confluence.ecmwf.int/display/CKB/ERA5%3A+data+documentation), which combines the model data with global climatic observations into to globally consistent dataset (Hersbach et al., 2018). CMIP6 includes more than 50 Global Circulation Models (GCMs), which are able to hindcast, as well as project climate for the next 100-200 years over the entire world. A set of emission scenarios driven by different socioeconomic assumptions, which are called “Shared Socioeconomic Pathways” (SSPs) has been developed to model the climate change outcomes. For the daily temperature data in 2020, we selected one of the GCMs (CNRM-CM6-1, France) which has a relatively higher spatial resolution and the highest emissions scenario (SSP5-8.5) for the temperature projections. We interpolated all the temperature data to a 1° × 1° resolution by using the Climate Data Operator (CDO; Schulzweida, 2019), then we calculated the seasonality and mean temperature (based on their definitions) for each grid cell in 2020 and 2080.

The bivariate plots (Fig. 4) of 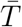 and *s* present the feasible global parameter space in 2020 and 2080. We treat the feasible parameter space in 2020 as the possible combinations of mean temperature and seasonality under current climate condition (Fig. 4a). Based on Fig. 4b, we see that the mean temperature in most regions will increase, while the seasonality may increase or decrease under future climate conditions. Note that the projections for 2080 suggest that we will see combinations of mean annual temperature and seasonality that are not currently present anywhere on the earth’s land surfaces (Fig. 4a). To simulate different climate change scenarios, we alter 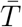 and *s* by adding 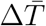 (0, 2, or 4) and Δs (−4, 0, or 4) across the entire parameter space (Fig. 4a), which generates nine different climate scenarios (one current climate scenario and eight future climate scenarios) (Fig. 5). We use these climate scenarios to study the role of biotic interactions in determining species population abundance under climate change.

**Figure 4:**
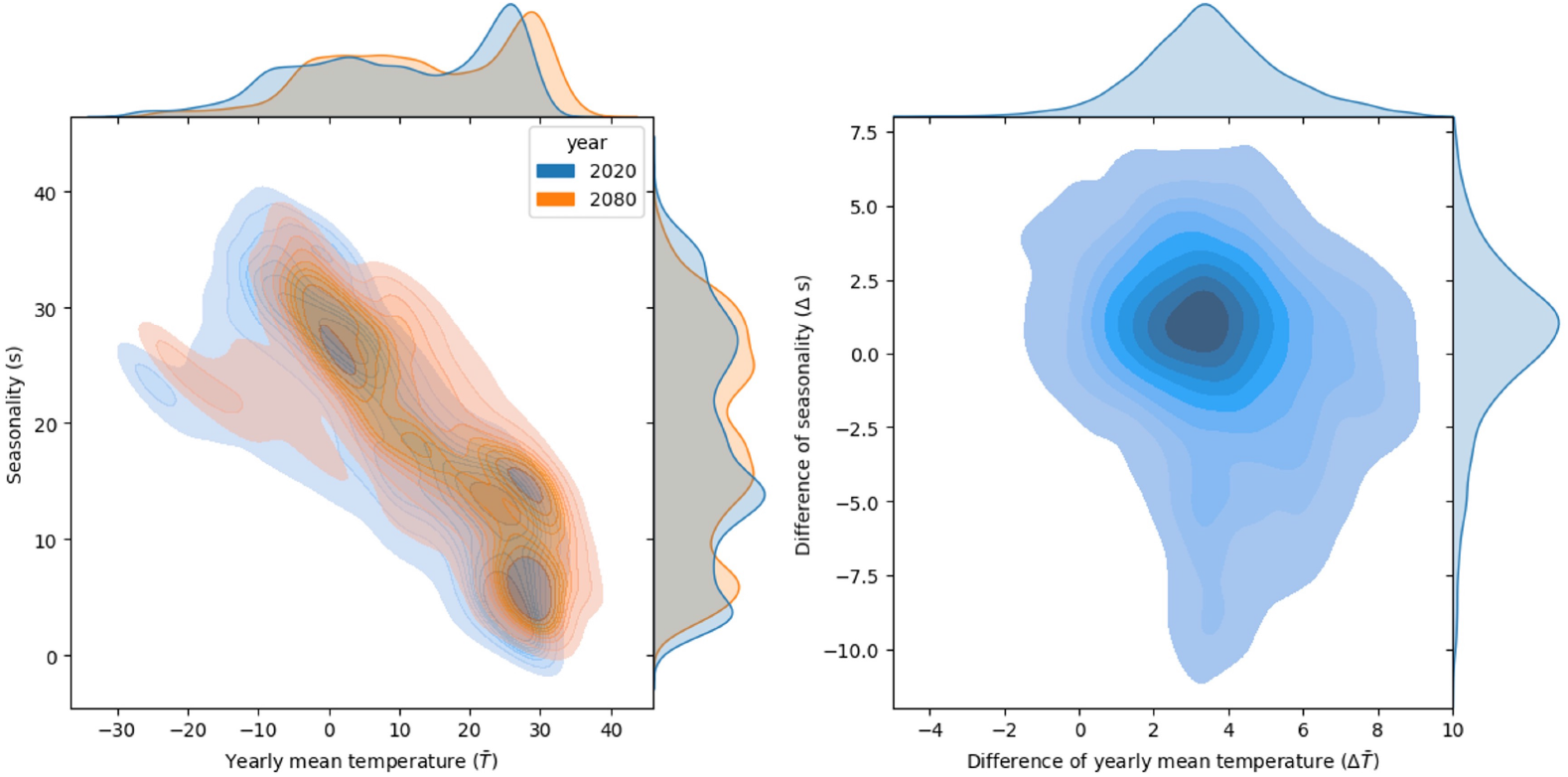
Feasible parameter spaces of yearly mean temperature 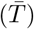 and seasonality (*s*) in 2020 and 2080. We calculated 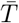 and *s* for all the grid cells across the entire land area (1° × 1° resolution) in 2020 and 2080 and plot the bivariate distributions (a) for 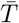 and *s* using kernel density estimation method. The univariate marginal distributions are also added along the x and y axes. (b) shows the bivariate distributions for the differences in 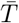 and *s* under climate change.

**Figure 5:**
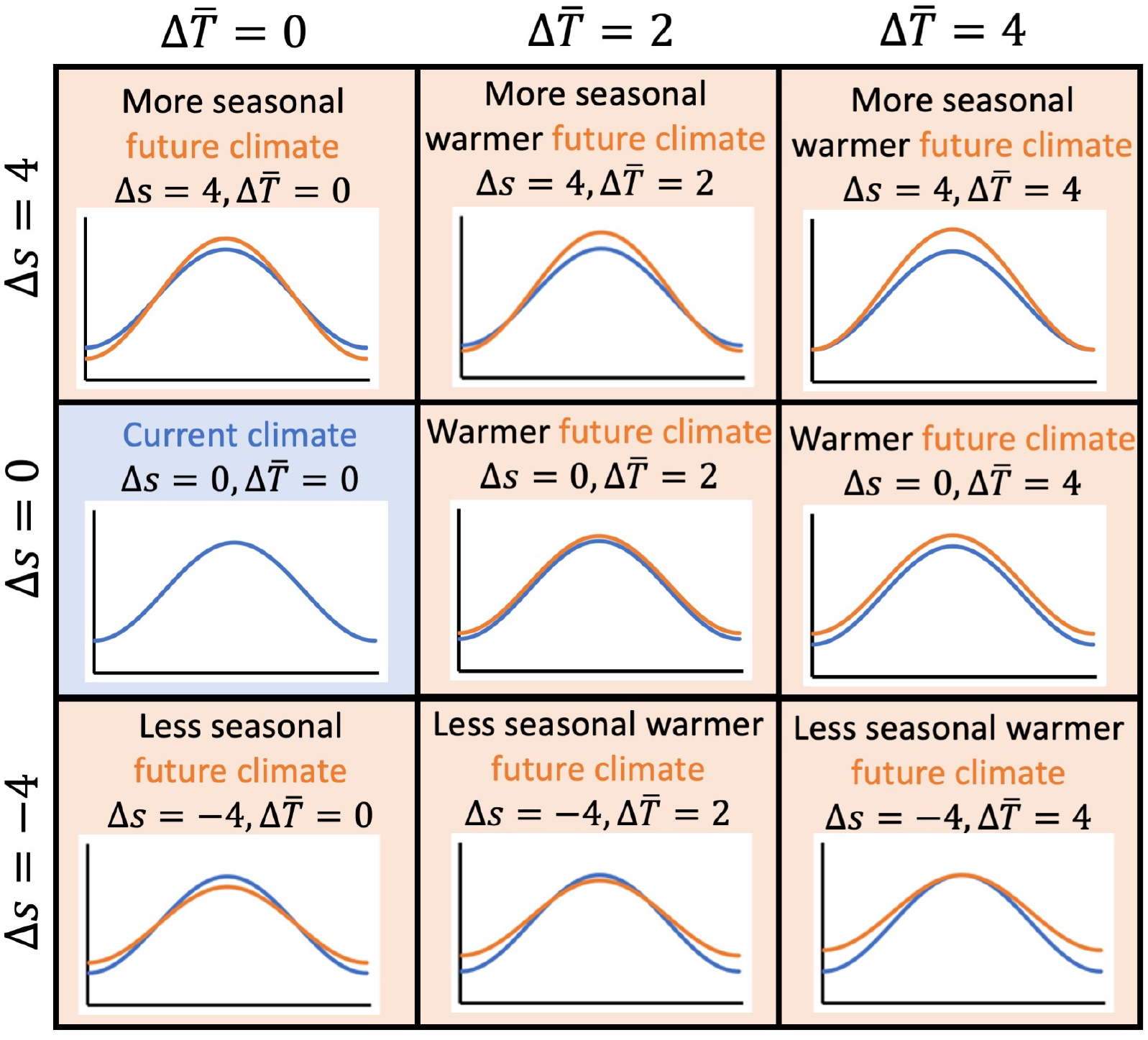
Conceptual diagrams for the nine climate scenarios. The eight future scenarios (with orange background) were generated by altering the yearly mean temperature 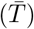 and seasonality (*s*). 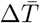 and Δ*s* represent the difference in 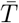 and *s* between different climate conditions (current and future).

### 2.4 Sensitivity analysis

To assess the robustness of our model and support the estimations of the parameters which lack of experimental data, we conducted a sensitivity analysis (SA) to evaluate the sensitivity of aphid and ladybird population growth, to changes in the aphid’s carrying capacity (*K*), the maximum intrinsic mortality rates for aphid (*υ_maxA_*) and ladybird (*υ_maxL_*), ladybird transformation rate (*Q_p_*), and the ratio of aphid’s initial abundance to the ladybird’s initial abundance (*R_initial_*).

For the purposes of the SA, we picked four points in each region of the parameter space. Each point have different temperature metrics (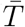 and *s*). We used Eq. 24 to model the daily mean temperatures throughout a year, which represent four temperature ‘profiles’ which result in different population abundance. The temperature metric values for the four temperature ‘profiles’ and the simulations of these temperature ‘profiles’ are shown in Fig. S2a1-d1.

The SA results provide guidance for uncertainty in these parameters. The results of the sensitivity analysis are shown in Fig. S2–S4. Except for *R_initial_*, our model results are not strongly affected by changing these parameters. The results for the *R_initial_* SA suggest the results fall into three regions. For low values of *R_initial_* (fewer aphids relative to ladybirds), ladybirds decimate the aphid population and then die out themselves from lack of prey. For high values of *R_initial_* (fewer predators relative to prey), rapid aphid reproduction allows the aphid population to ‘escape regulation’ by the predator (Fig. S4) and thereafter are *regulated by their carrying capacity* (*K*). These two solution spaces are uninteresting as our focus was on the importance of the predator-prey interaction, which is largely irrelevant in these regions of parameter space. For intermediate values of *R_initial_*, prey populations are mostly *regulated by the predators*. The exact ratios that define the boundaries of these three regions of the solution space depend on the particular values of 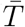 and *s*. As our focus was on the predator-prey interaction under climate change, we choose an intermediate value of *R_initial_* for the main simulations for this study. Note as well that the results are more sensitive to *υ_maxL_* than *υ_maxA_* (Fig. S5 and S6). Among the different climate conditions, species that live in warmer less seasonal climate are more sensitive to these two parameters.

### 2.5. Model analysis

In the feasible parameter space under current climate (Fig. 4a), we uniformly sampled 290 combinations of the two temperature metrics (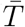 and *s*). Each sampled combination of 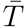 and *s* determine the daily temperatures throughout of a year as defined by Eq. 24.

Each simulation began with 1 × 10^5^ aphids and 500 ladybird beetles (per 100 *m*^2^, split equally among all stages; *R_initial_* = 200) introduced separately on the dates for which temperatures were ‘warm enough’ (*T* > *CT_min_*) to support aphids’ and ladybird beetles’ positive growth. If predators are more cold tolerant than prey, we do not introduce ladybirds until the temperatures reach to aphid’s lower temperature threshold. Accordingly, predators are less cold tolerant than prey, we introduce the aphids ahead of ladybirds. This approach may be biologically unrealistic. Phenological mismatches will happen under climate change (see e.g., Visser and Gienapp, 2019) and this is an import impact of climate change. However, as our focus was on the predator-prey interactions *per se*, we chose an approach that eliminated such mismatches, and assumed ladybirds time emergence when positive growth is supported. Each simulation was terminated once the aphid and ladybird’s daily population abundance were both < 1, or when the end of the simulation year was reached.

For each pair of interacting species, we ran the simulations for all the temperature profiles under nine different climate scenarios (Fig. 5) and obtained the aphid and ladybird population abundances (*AAP* and *ALP*) across the feasible parameter space under different climate scenarios. Then, we compared species abundances under climate change among the nine pairs of species which have different thermal tolerances, to gain insight into the generalized responses of a predator-prey system to climate change.

## 3. Results

### 3.1. Result classes

Heatmaps of abundance generally exhibit three or four distinct regions. The *relative* magnitudes of abundances in these four regions fall into one of three classes, which we denote as ‘*A*-Class’, ‘*B*-Class’ and ‘*C*-Class’. These are illustrated in Fig. 6. The location of the regional boundaries change in different heatmaps, but the magnitude of values in different regions are roughly consistent for each class. In the *A*-Class, the relative magnitudes are *A*_1_ = 0 < *A*_2_ ≈ *A*_4_ < *A*_3_. In the *B*-Class, the relative magnitudes are *B*_1_ = 0 < *B*_3_ < *B*_2_ ≈ *B*_4_. In the ‘*C*-Class, the relative magnitudes are *C*_1_ = 0 < *C*_2_ < *C*_3_ < *C*_4_. Note further, it is helpful to link these regions in terms of combinations of 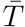 and *s*. For example, region 3 (*A*_3_, *B*_3_ and *C*_3_) corresponds to warmer less seasonal regions, and there can be a qualitative shift in species abundance moving from *A*-class to *B*-class to *C*-class.

**Figure 6:**
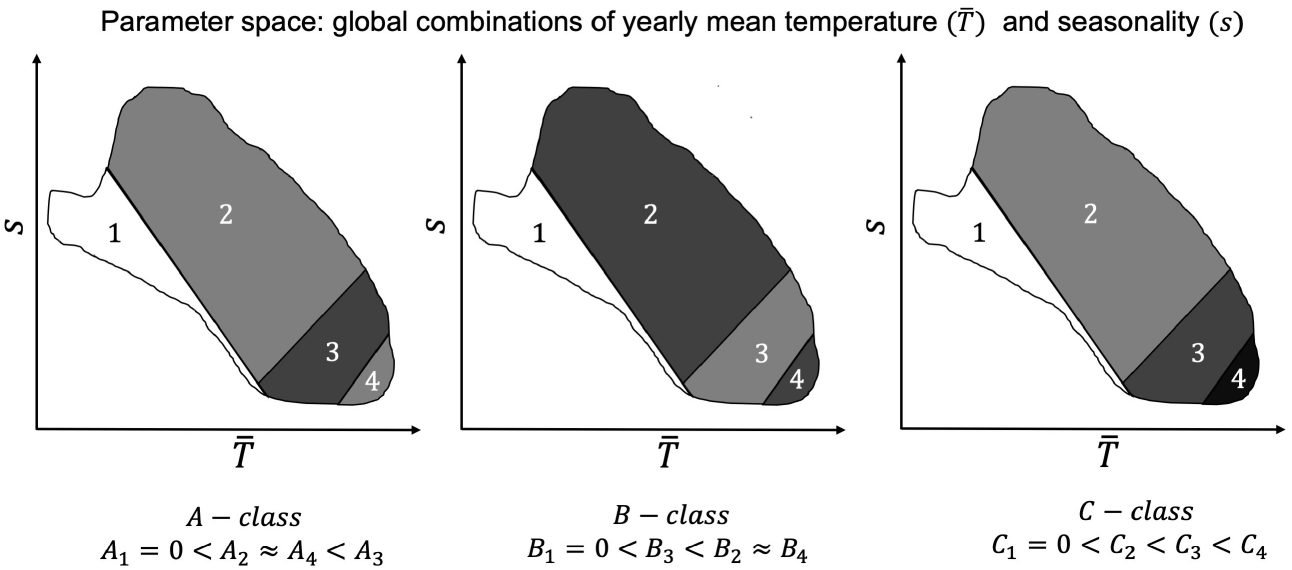
Result classes in parameter space. The parameter space represents all the possible combinations of mean temperature 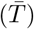 and seasonality (*s*). In the *A*_1_, *B*_1_, *C*_1_-regions, the annual abundances is 0. This region of parameter space is too cold for persistence. Note that the region 4 is not always present in A-Class results. The exact boundaries between each region in each class depend on the particular climate scenario and the relationships among the critical temperature thresholds. The ordinal patterns are the defining feature.

### 3.2. *Identical thermal niches* (AL1: base case)

Fig. 7a shows *AAP* across the feasible parameter space when prey and predator have identical thermal niches for predators and prey. These are *A*-Class results. The largest *AAP* values occur in the *A*_3_-region (see Fig. 6). We see that the *A*_3_-region of parameter space increases as the climate becomes warmer (larger values of 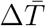) and less seasonal (smaller values of Δ*s*). The *A*_3_-region expands towards the colder, less seasonal regions. This makes sense intuitively since this area of the parameter space is becoming more similar to the *A*_3_-region under the current climate (Fig. 7a4). In Figures 7a2 and 7a3, we see the appearance of the *A*_4_-region. Here, the warmest and least seasonal region of the parameter space is becoming too warm for the aphids (note the change of color from green to blue). Notice as well that the size of the *A*_1_-region decreases as the the climate becomes warmer (larger values of 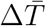) and increases as the climate becomes more seasonal (larger values of Δ*s*). As the climate warms, more of the region that was formerly too cold for aphid persistence (*A*_1_-region) becomes suitable. Solutions along the *A*_1_-*A*_2_ boundary become unsuitable for the aphids (A_1_-region) as the climate becomes less seasonal (lower Δ*s*). In other words, conditions along this boundary are becoming ‘more constantly cold’ and hence unsuitable for aphid development.

**Figure 7:**
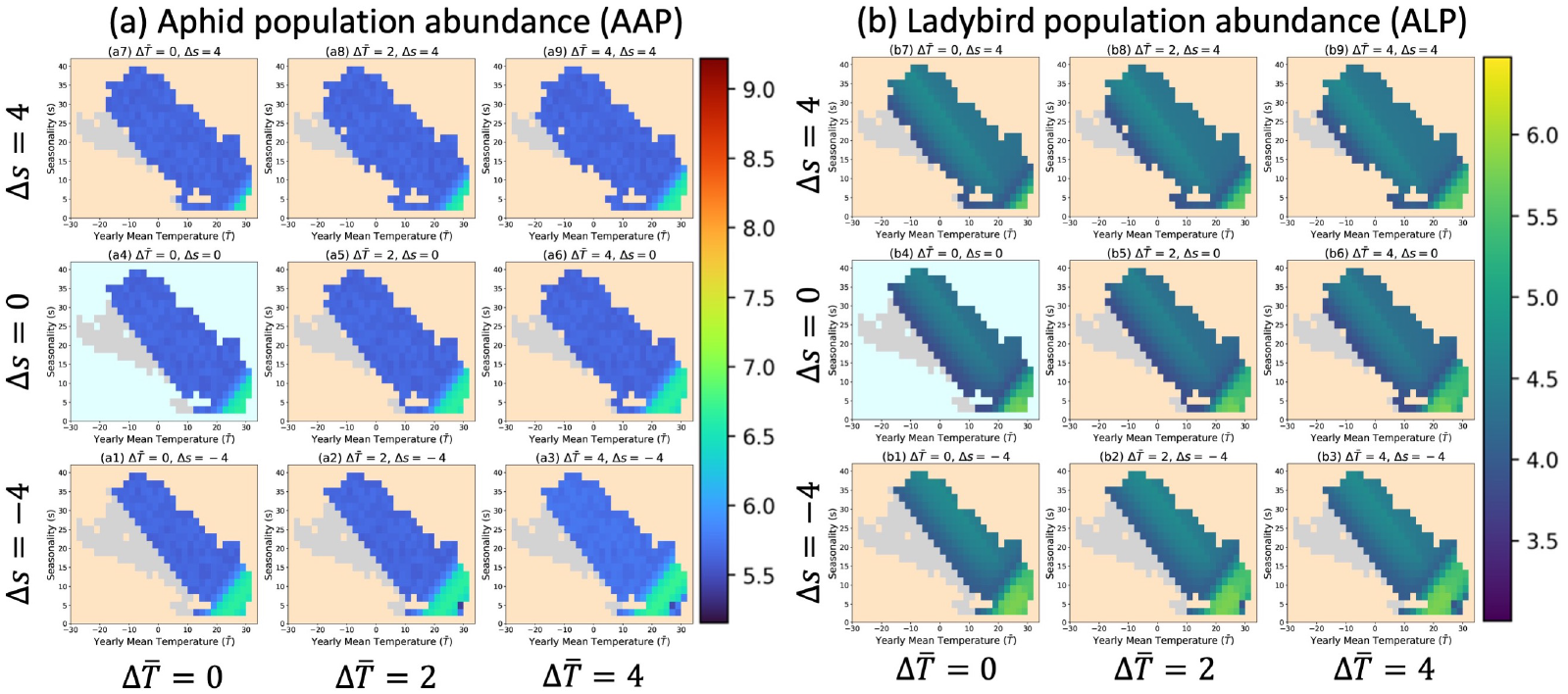
Heatmaps in parameter spaces for aphid and ladybird population abundance under different climate scenarios (*AL1: identical thermal niches*, *A*-Class). The left nine heatmaps (a) represent the patterns for aphid abundance. The right nine heatmaps (b) represent the patterns for ladybird abundance. Colored region represents the global combinations for mean temperature and seasonality. The points with light grey color indicate that climates are unsuitable for the prey or predator. The heatmap with light blue background represents the abundance pattern under current climate, the rest heatmaps with light orange backgrounds represent the abundance patterns under different future climates.

Fig. 7b shows the same information for the ladybird abundance (*ALP*). It shows the same pattern as the *AAP* results and, not surprisingly, *ALP* is maximal when *AAP* is maximal because there are more prey available to fuel ladybird population growth.

### 3.3. *Predators have* wider *thermal niche breadths than the prey*

#### *Predators are more cold tolerant* (AL2)

The heatmaps (Fig. S7) show the same general patterns (*A*-Class) as the base case (AL1, Fig. 7) except that prey and predators are less abundant than in the base case. More cold tolerant predators are, *ceterius parabis*, better able to control the prey. For more detailed analyses of this and the following species pairs, see the supplemental information.

#### *Predators are more heat tolerant* (AL3)

Fig. S8) again shows the same general pattern (*A*-Class) as the base case (AL1, Fig. 7), but the population abundance for both species are slightly smaller than AL1, especially in *A*_3_ and *A*_4_ regions.

#### *Predators are more heat and more cold tolerant* (AL4)

Heatmaps for AL4 (Fig. S9) show a very similar pattern to the base case (Fig. 7a) as well as the AL2 and AL3. The prey abundance is more similar to AL3 than the other two pairs.

### 3.4. *Predators have a* narrower *thermal niche breadth*

#### *Predators are less cold tolerant* (AL5)

We see a somewhat different pattern (Fig. 8) than we see in the base case (Fig. 7) or when predators have a wider thermal niche than the prey (AL2–AL4). Here, the results are now of the *B*-Class (see Fig. 6). Unlike previous pairs which have much higher population abundance in *A*_3_-regions, predators and prey are less abundant in the *B*_3_-regions than *B*_2_-regions. Also, the *B*_3_-regions are larger under warmer less seasonal climate conditions. The other obvious difference with the previous pairs is the much greater aphid abundance in the *B*_2_-region compared with the *A*_2_-regions in those cases. With a narrower thermal niche breadth, the predator does worse, *ceterius parabis*, and so prey are less well controlled. Nevertheless, greater prey abundance leads to greater predator abundance. The predator abundance differences between *B*_2_ and *B*_3_ regions are not as large as those for the prey abundance.

**Figure 8:**
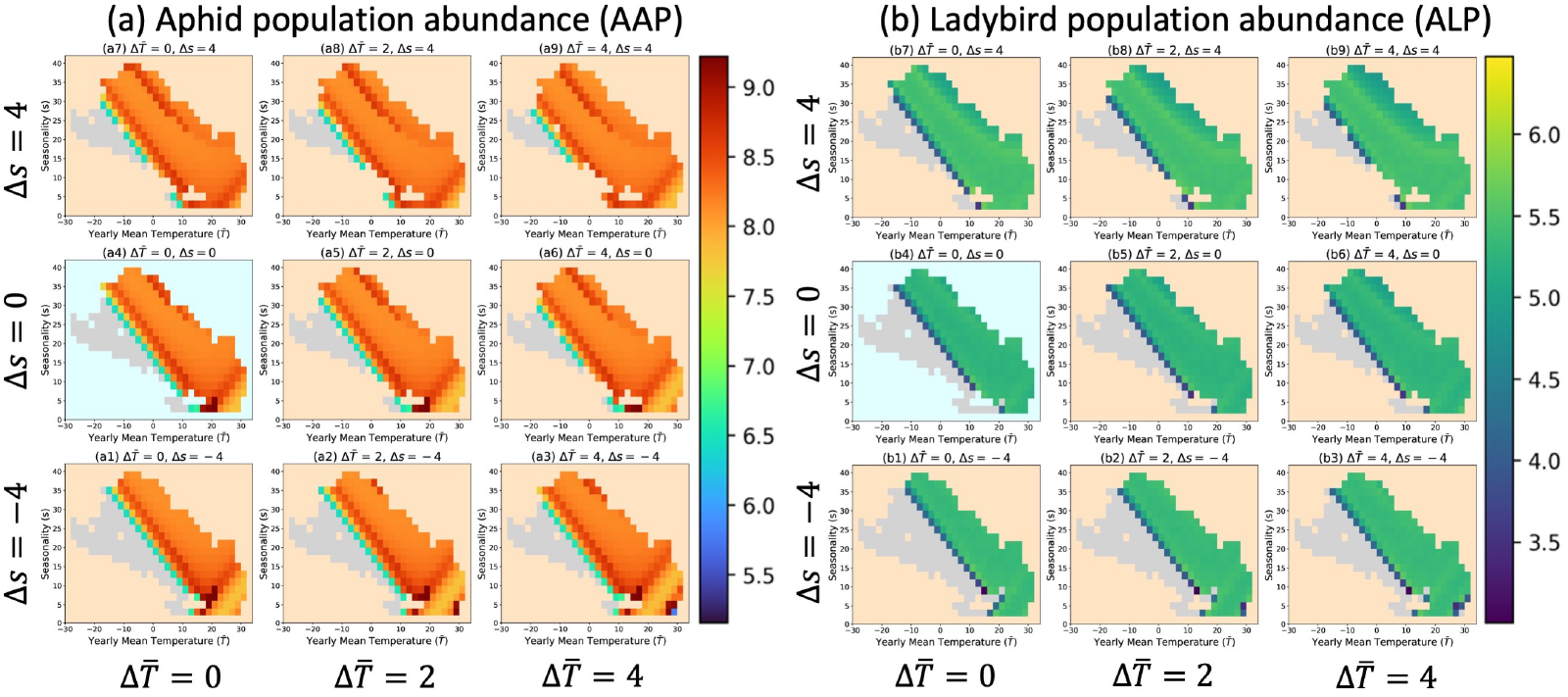
Heatmaps in parameter spaces for aphid and ladybird population abundance under different climate scenarios (*AL5: predators are less cold tolerant than the prey*, *B*-Class). Colored region represents the global combinations for mean temperature and seasonality. The left nine heatmaps (a) represent the patterns for aphid abundance. The right nine heatmaps (b) represent the patterns for ladybird abundance. The points with light grey color indicate that climates are unsuitable for the prey or predator. The heatmap with light blue background represents the abundance pattern under current climate, the rest heatmaps with light orange backgrounds represent the abundance patterns under different future climates.

#### *Predators are less heat tolerant* (AL6)

The patterns for aphid abundance (Fig. 9a) are *C*-Class, which are similar to the base case condition (Fig. 7) but differ in regions 3 and 4. Aphids are much more abundant in the *C*_3_-region here than in the previous cases we examined. With warming climate, aphid abundance is greater for the warmest and least seasonal climates (*A*_4_-region). This occurs because under these conditions ‘more constantly hot’ climate limits the development of ladybirds when predators are less heat tolerant, allowing aphid populations to grow larger.

**Figure 9:**
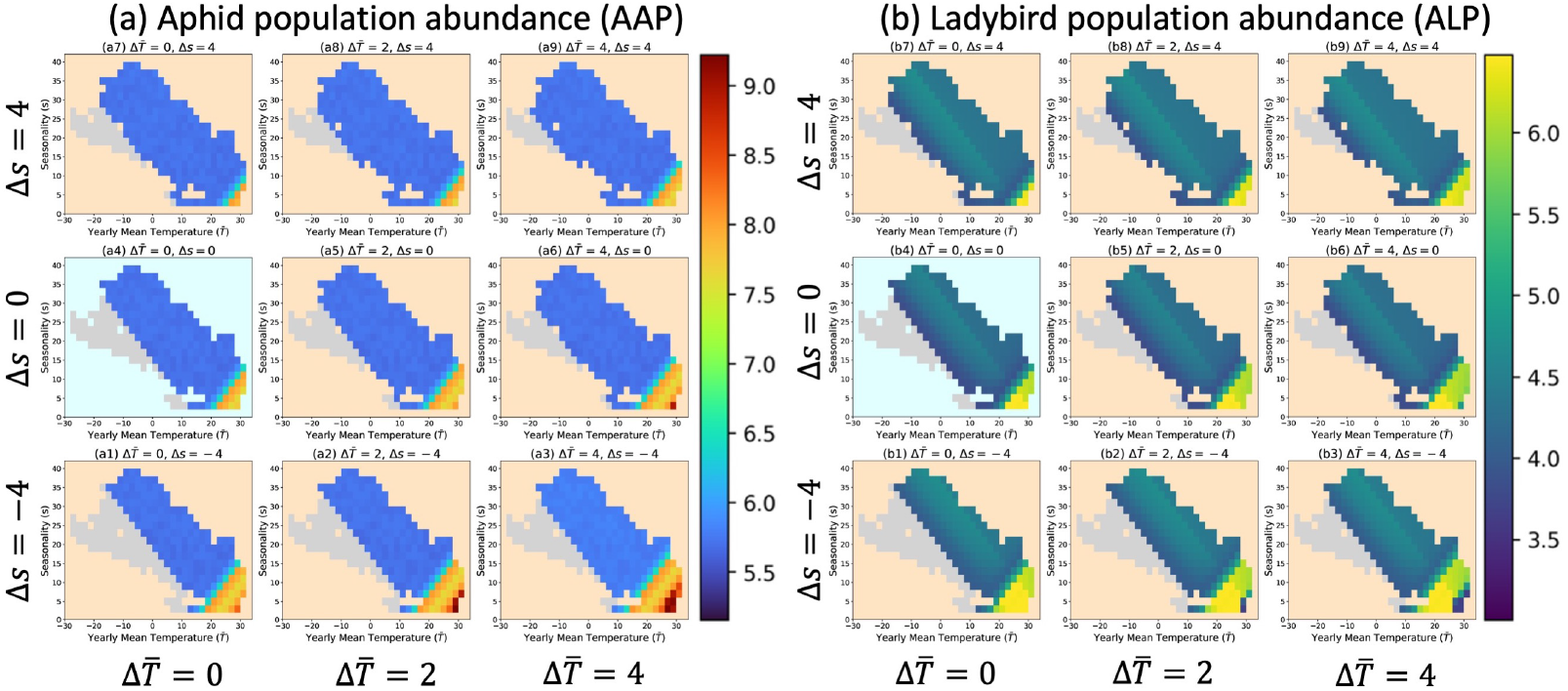
Heatmaps in parameter spaces for aphid and ladybird population abundance under different climate scenarios (*AL6: predators are less heat tolerant*, *C*-Class). Colored region represents the global combinations for mean temperature and seasonality. The left nine heatmaps (a) represent the patterns for aphid abundance. The right nine heatmaps (b) represent the patterns for ladybird abundance. The points with light grey color indicate that climates are unsuitable for the prey or predator. The heatmap with light blue background represents the abundance pattern under current climate, the rest heatmaps with light orange backgrounds represent the abundance patterns under different future climates.

#### *Predators are both less cold and less heat tolerant* (AL7)

Fig. S10 is quite similar to AL5 and is *B*-Class. Again, as in AL2, aphid abundance is quite high compared to when predators have a wider thermal niche than the prey.

Notice that the *A*-Class results change to *B*-Class when the predators are less cold tolerant than the prey (AL5 and AL7).

### 3.5. Identical thermal breadths

#### *Predators are more cold and less heat tolerant than the prey* (AL8)

Fig. S11a is a *C*-Class result. In general, the prey abundance is lower than AL7 and the base case but a bit higher than AL2. This matches our intuition in that the lower heat tolerance of the predators offsets, in part, more cold tolerance in decreasing prey abundance.

#### *Predators are less cold and more heat tolerant than the prey* (AL9)

Just as in the other pairs of species where the predator is less cold tolerant than the prey (AL5 and AL7), Fig. S12a shows *B*-Class results. These results are also similar in magnitude to those cases.

### 3.6. Summary for all species pairs

Comparing all the heatmaps for each pair (Fig. 10), cold tolerance of predators is much more influential than heat tolerance on prey abundance. Furthermore, when the predator has a wider thermal tolerance niche, it is most able to control the prey population; in contrast, when the predator has a narrower thermal tolerance niche, it poorly controls the prey population. Overall, the mismatch of minimum and maximum thermal tolerances between predators and prey matters in determining species’ response to the changing climate, even when both species have identical thermal tolerance breadth.

**Figure 10:**
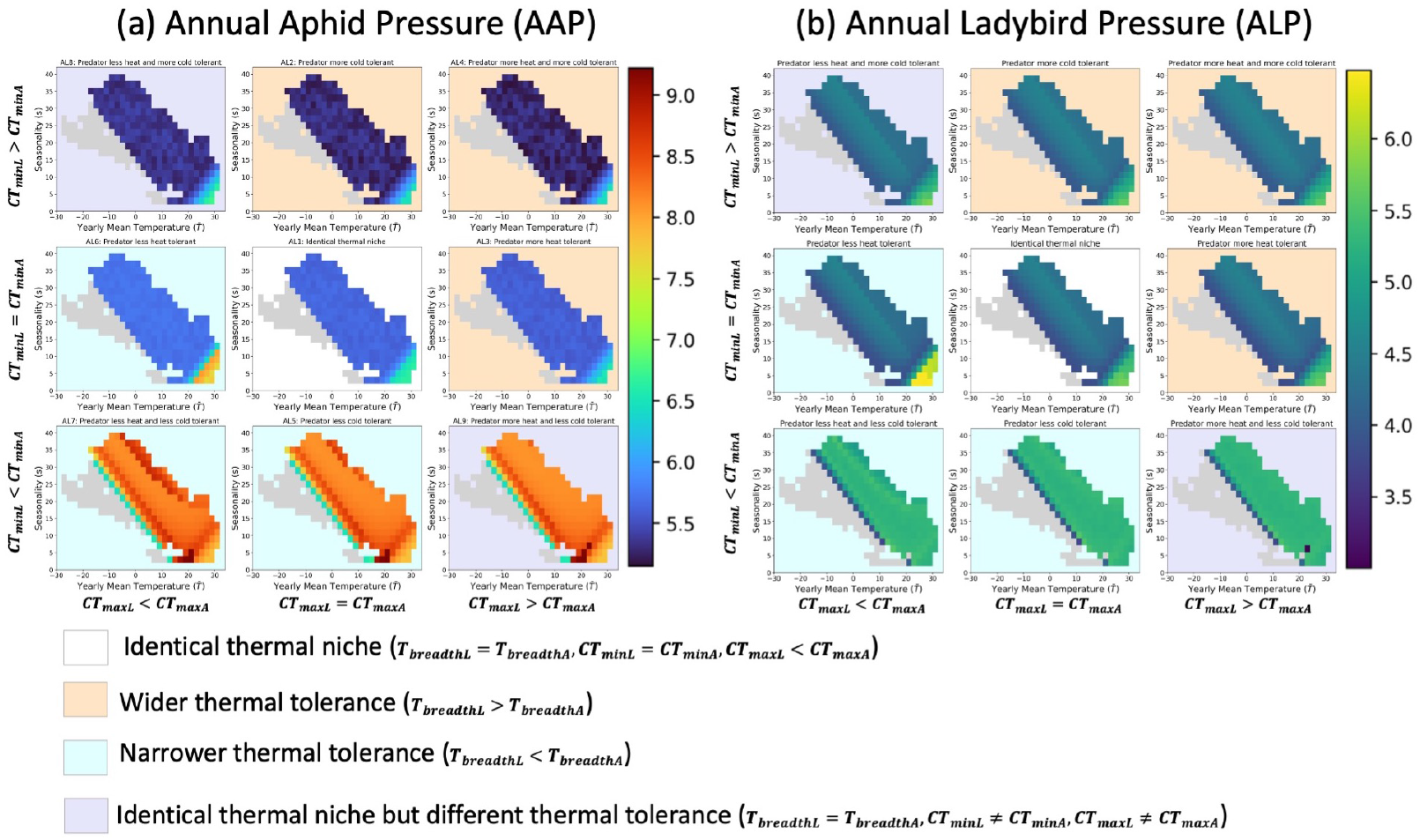
Heatmaps for nine thermal tolerance mismatching predator-prey pairs under current climate. (a1)–(a9) and (b1)–(b9) represent the heatmaps for aphid and ladybird population abundance, respectively. The heatmap with white background represent the species pairs with identical thermal niche, three heatmaps with light orange background represent the species pairs that predators have wider thermal tolerances, the three heatmaps with light blue background represent the species pairs that predators have narrower thermal toleranes, and the two heatmaps with light purple.

In response to climate change, all scenarios have a coherent response in the light grey region (*A*_1_, *B*_1_ and *C*_1_ regions), i.e., a warmer more seasonal future climate will lead more regions becoming suitable for these species to survive. In the regions with warmer less seasonal climates (*A*_3_ and *B*_3_ regions), predators and prey with different thermal tolerances will respond to climate change differently. Except for the three pairs when predators are less cold tolerant than the prey (AL5, AL7, AL9), prey will be more abundant when future climates become warmer and less seasonal in *A*_3_ region. However, prey will be less abundant in *B*_3_ region under such a climate change scenario. Notably, there is a trade-off between warmer climate and more seasonal climate, which makes it difficult to predict the effect of a more seasonal climate on species abundance.

## 4. Discussion

It is traditional to construct dynamic stage-structured models of insect populations. To date, a large variety of such models have been developed for studying predator-prey interactions (see e.g., Kettle and Nutter, 2015; Kha-janchi, 2014; Wang and Zou, 2017; Lu et al., 2017; Xia et al., 2018; Mortoja et al.,2018) as well as herbivore-plant interactions (see e.g., Newman et al., 2003; Newman, 2004, 2005, 2006; Thornley and Newman, 2022). These studies have a variety of aims such as model improvement, biological control and in-season forecasting. For example, Kettle and Nutter (2015) present an R-package, stagePop, which can be used to simulate the deterministic dynamics of stage-structured populations that are involved in species interactions, environmental change and so on. They have taken the Beddington–DeAngelis-type functional response (Beddington, 1975; DeAngelis et al., 1975), which admits rich and biologically meaningful dynamics to study predator-prey interactions. Xia et al. (2018) developed a detailed process-based simulation model which includes interactions and is affected by temperature and host growth for the biological control of the cotton aphid. All of these studies provide insights into predator-prey interactions. Here, we sought to understand how thermal tolerance mismatches affect predator-prey interactions under climate change.

The use of stage-structured models to study the impacts of climatic change in combination with thermal performance are few. Skirvin et al. (1997) modified their earlier model of the population dynamics of *Sitobion avenae* to incorporate a ladybird beetle predator *Coccinella septempunctata*, and then predicted the likely effects of climatic change on their interactions. They found that coccinellids are most effective at reducing aphid abundance in relatively hot summers. Hoover and Newman (2004) developed a mechanistic model of a tri-trophic interaction between grass, cereal aphids (*Rhopalosiphum padi*) and their parasitoids (*Aphidius rhopalosiphi*) and examined the interacting effects under climate change. Their model predicted that parasitoids do not fundamentally alter the aphid response to climate change. Although these studies have a similar focus to our study, they are specific to particular pairs of interacting species, and neither considered the heterogeneity of interacting species’ thermal tolerance mismatch.

Thermal performance mismatches among interacting species are common in nature (Agosta et al., 2018). Laboratory studies on the thermal limits of *Aphis gossypii* and its predator *Harmonia dimidiata*, showed that the aphid has a higher *CT_max_* than the ladybird (Yu et al., 2013). Hughes et al. (2010) found that the parasitoid *Lysiphlebus testaceipes* is more cold and more heat tolerant than its host, the black bean aphid (*Aphis fabae*). Agosta et al. (2018) found that the caterpillar host *Manduca sexta* had a higher *CT_max_* (≈ 4°C) than the parasitic wasp, *Cotesia congregata*, and the hyperparasitic wasp, *Conura* sp., had a higher *CT_max_* (≈ 6°C) than its host, *C. congregata*. Pintanel et al. (2021) studied a predator-prey system of dragonflies (20 species) and anuran larvae (17 species). Their analyses revealed that predators exhibit higher heat tolerances than prey (≈ 4°C) across habitats and elevations. Buxton et al. (2020) examined the thermal tolerance of three mosquito species and their predators and concluded that the predators had lower *CT_min_* and *CT_max_* than the mosquito prey. These inconsistent thermal tolerance performances among different predator and prey species point to the significance of our generalized approach.

We comprehensively examined the generalized responses of a predator-prey system to climate change by using pairs of interacting species with different thermal tolerances. Our results show prey abundance is affected by the predator’s thermal tolerance. When the predators’ thermal tolerance is narrower than that of the prey, prey abundance increases, especially when predators are less cold tolerant than the prey. Cold tolerance of the predator is much more influential than their heat tolerance on prey abundance. Poorer cold tolerance doesn’t allow the predator to regulate prey growth at the beginning of the growing season. Notably, Bennett et al. (2021) have concluded that cold tolerance has evolved more quickly than heat tolerance in both endotherms and ectotherms, which indicates that the evolutionary adaption of a species’ cold tolerance may lead to significant impacts on interacting species. Our results also suggest the importance of integrating evolutionary adaption into the study of species interactions.

We have shown that the outcome of the predator-prey interaction depends on the relationship between the mean annual temperature and the seasonality. Climate change projections indicate warming, to varying degrees, all over the globe, and generally an increase in seasonality, except at high latitudes and altitudes (Fig. S13). Currently, aphids and ladybirds do well in tropical regions where the mean annual temperature is between the *CT_max_* and *CT_min_* values for each species and low seasonality ensure that both species spend most or all of the year in temperatures conducive to growth. As climate change pushes the mean temperature above the *CT_max_* and increases seasonality, some or all of the year will become too warm to support positive growth rates. Conversely, at high latitudes, the current mean annual temperature is below the *CT_min_* values, the low seasonality means that the temperature is rarely, if ever, above *CT_min_*. As the climate warms and becomes more seasonal, some portion of the year may become warm enough to support positive population growth rates. In the middle latitudes, we see both decreases and increases in the amounts of time when the temperature is between the *CT_max_* and *CT_min_* values. In the lower middle latitudes the increasing temperature and seasonality tend to make too warm parts of the year that were previously suitable. The reverse is true for the upper middle latitudes where increasing temperature and seasonality tend to make more of the year suitable (i.e., < *CT_max_* and > *CT_min_*). Our findings are consistent with latitudinal patterns in Youngsteadt et al. (2017)’s experimental study, which shows that species abundance will increase with warming in high-latitude taxa, but have heterogeneous responses in mid-latitudes.

The lack of evolution in the species’ thermal tolerance performances in response to climate change is a significant limitation of our study. Predators and prey may evolve differently in respond to environmental change, which will alter their subsequent interactions (Grigaltchik et al., 2012; Cheng et al., 2017). The shapes of predator’s and prey’s thermal performance curves may shift in the face of climate change. Tüzün and Stoks (2018) considered six scenarios of how an evolutionary shift in the thermal tolerance of one species may affect the performance of its interacting species and concluded that evolution of thermal tolerance curves may strongly impact the predicted outcome of biotic interactions under climate warming. However, they did not support their analysis with general theory nor systematically predict the outcomes. In Tüzün and Stoks (2018)’s study, the three scenarios that involve a horizontal shift in thermal tolerance are analogous to the comparisons in our study between AL1 and AL9, AL2 and AL3, and AL6 and AL5. Our modelling leads to the conclusion that for both prey and predator abundances will increase if the predator’s thermal performance curve shifts to the right horizontally, which supports Tüzün and Stoks’ (2018) conclusion about the importance of considering the evolution of species thermal tolerances. The abiotic environment and species interactions may drive the natural selection of species’ thermal limits in the long term. For example, when predators have narrower thermal niches, it eventually leads to higher abundances for both prey and predators, but it is unclear how thermal performance differences lead to differences in individual fitness. The evolution of thermal performance of interacting species to climate change adds uncertainties to the prediction of species’ abundances. Prey may respond faster to climate change than predators due to a shorter generation time but the long periods during which aphids reproduce asexually may mean that they evolve more slowly than their predators. We will examine the influence of evolution on predator and prey thermal tolerance in a future paper.

Although the current model might be seen as ‘biologically detailed’, there are still some limitations due to the trade-off between simplicity and realism (Levins, 1966). We made assumptions and model choices for simplicity and tractability. For example, we only modeled growing season abundances and assumed the aphid and ladybird population are closed (i.e., no immigration and emigration), as well as ignored other sources of species’ extrinsic mortality rate. Furthermore, prey abundances are highly dependent on the initial prey-predator ratio (*R_initial_*), which is a key factor for biological control, and also supported by Latham and Mills’ (2010) and Xia et al.’s (2018) studies. Nevertheless, when predators are less cold tolerant than the prey, the predators are not able to regulate the prey population at the beginning of the growing season because they develop slower and later than the prey. Predators were not able to regulate the prey population no matter how low we set the *R_initial_*. Although the patterns of species abundance are different with different *R_initial_* (Fig. S14), general conclusions stay the same no matter what the early season densities for predators and prey are.

## 5. Conclusions

We used the biologically detailed stage-structured population dynamic model present in this study to summarize the generalized response of a predator-prey system with different thermal tolerances to climate change. Notwithstanding the limitations and uncertainties, our study identify three common patterns of species abundance across the feasible parameter space that relate to the type of thermal tolerance mismatches. Our results indicate that thermal tolerance mismatch between predators and prey affects their abundance, as well as their response to climate change. The main findings of this study suggest the dominant role of cold tolerance in affecting prey abundance, especially under climate change scenarios. Our study highlights the significance of understanding how thermal tolerance mismatches affect species interactions.

## 6. Author contribution

**Xuezhen Ge**: Conceptualization, Methodology, Project administration, Software, Visualization, Writing - original draft, Writing - review and editing. **Cortland K. Griswold**: Conceptualization, Funding acquisition, Methodology, Writing - review and editing. **Jonathan A. Newman**: Conceptualization, Funding acquisition, Methodology, Writing - review and editing.

## 7. Acknowledgements

We thanks Sharcnet and Compute Canada for providing computational support. This work was conducted on the traditional territory of the Neutral, Anishnaabe and Haudenosaunee peoples. It is important to acknowledge these peoples as the traditional stewards of the land and doing so reminds us of our on-going responsibilities toward reconciliation.

## References

Agarwala, B.K., Singh, T.K., Lokeshwari, R.K., Sharmila, M., 2009. Functional response and reproductive attributes of the aphidophagous ladybird beetle, *Harmonia dimidiata* (Fabricius) in oak trees of sericultural importance. Journal of Asia-Pacific Entomology 12, 179–182.

Agosta, S.J., Joshi, K.A., Kester, K.M., 2018. Upper thermal limits differ among and within component species in a tritrophic host-parasitoid-hyperparasitoid system. PLoS One 13, e0198803.

Aldyhim, Y., Khalil, A., 1993. Influence of temperature and daylength on population development of aphis gossypii on cucurbita pepo. Entomologia Experimentalis et Applicata 67, 167–172.

Alexander, J.M., Diez, J.M., Levine, J.M., 2015. Novel competitors shape species’ responses to climate change. Nature 525, 515–518.

Amundrud, S.L., Srivastava, D.S., 2020. Thermal tolerances and species interactions determine the elevational distributions of insects. Global Ecology and Biogeography 29, 1315–1327.

Beddington, J.R., 1975. Mutual interference between parasites or predators and its effect on searching efficiency. The Journal of Animal Ecology, 331–340.

Bennett, J.M., Sunday, J., Calosi, P., Villalobos, F., Martínez, B., Molina-Venegas, R., Araújo, M.B., Algar, A.C., Clusella-Trullas, S., Hawkins, B.A., et al., 2021. The evolution of critical thermal limits of life on earth. Nature communications 12, 1–9.

Bianchi, F.J., Booij, C., Tscharntke, T., 2006. Sustainable pest regulation in agricultural landscapes: a review on landscape composition, biodiversity and natural pest control. Proceedings of the Royal Society B: Biological Sciences 273, 1715–1727.

Birkett, A.J., Blackburn, G.A., Menéndez, R., 2018. Linking species thermal tolerance to elevational range shifts in upland dung beetles. Ecography 41, 1510–1519.

Blois, J.L., Zarnetske, P.L., Fitzpatrick, M.C., Finnegan, S., 2013. Climate change and the past, present, and future of biotic interactions. Science 341, 499–504.

Boukal, D.S., Bideault, A., Carreira, B.M., Sentis, A., 2019. Species interactions under climate change: connecting kinetic effects of temperature on individuals to community dynamics. Current opinion in insect science 35, 88–95.

Brown, M., 2004. Role of aphid predator guild in controlling spirea aphid populations on apple in West Virginia, USA. Biological Control 29, 189–198.

Buxton, M., Nyamukondiwa, C., Dalu, T., Cuthbert, R.N., Wasserman, R.J., 2020. Implications of increasing temperature stress for predatory biocontrol of vector mosquitoes. Parasites & Vectors 13, 1–11.

Chen, I.C., Hill, J.K., Ohlemüller, R., Roy, D.B., Thomas, C.D., 2011. Rapid range shifts of species associated with high levels of climate warming. Science 333, 1024–1026.

Cheng, B.S., Komoroske, L.M., Grosholz, E.D., 2017. Trophic sensitivity of invasive predator and native prey interactions: integrating environmental context and climate change. Functional Ecology 31, 642–652.

Damien, M., Tougeron, K., 2019. Prey–predator phenological mismatch under climate change. Current opinion in insect science 35, 60–68.

DeAngelis, D.L., Goldstein, R., O’Neill, R.V., 1975. A model for tropic interaction. Ecology 56, 881–892.

Dedryver, C.A., Le Ralec, A., Fabre, F., 2010. The conflicting relationships between aphids and men: a review of aphid damage and control strategies. Comptes Rendus Biologies 333, 539–553.

Farhadi, R., Allahyari, H., Chi, H., 2011. Life table and predation capacity of *Hippodamia variegata* (Coleoptera: Coccinellidae) feeding on *Aphis fabae*(Hemiptera: Aphididae). Biological Control 59, 83–89.

Gilman, S.E., 2017. Predicting indirect effects of predator–prey interactions. Integrative and comparative biology 57, 148–158.

Glen, A.S., Dickman, C.R., 2014. The importance of predators. Carnivores of Australia: past, present and future. Australia: CSIRO Publishing, 1–12.

Grigaltchik, V.S., Ward, A.J., Seebacher, F., 2012. Thermal acclimation of interactions: differential responses to temperature change alter predator–prey relationship. Proceedings of the Royal Society B: Biological Sciences 279, 4058–4064.

Hersbach, H., Bell, B., Berrisford, P., Biavati, G. and Horányi, A., Muñoz Sabater, J., Nicolas, J., Peubey, C., Radu, R., Rozum, I., et al., 2018. ERA5 hourly data on single levels from 1979 to present. https://cds.climate.copernicus.eu/cdsapp#!/dataset/reanalysis-era5-single-levels?tab=overview. doi:10.24381/cds.adbb2d47. date accessed: December 14, 2019.

Holling, C.S., 1959. Some characteristics of simple types of predation and parasitism. Canadian entomologist 91, 385–398.

Hoover, J.K., Newman, J.A., 2004. Tritrophic interactions in the context of climate change: a model of grasses, cereal aphids and their parasitoids. Global Change Biology 10, 1197–1208.

Hughes, G.E., Owen, E., Sterk, G., Bale, J.S., 2010. Thermal activity thresholds of the parasitic wasp lysiphlebus testaceipes and its aphid prey: implications for the efficacy of biological control. Physiological Entomology 35, 373–378.

Kersting, U., Satar, S., Uygun, N., 1999. Effect of temperature on development rate and fecundity of apterous aphis gossypii glover (hom., aphididae) reared on gossypium hirsutum l. Journal of Applied Entomology 123, 23–27.

Kettle, H., Nutter, D., 2015. stagepop: Modelling stage-structured populations in r. Methods in Ecology and Evolution 6, 1484–1490.

Khajanchi, S., 2014. Dynamic behavior of a beddington–deangelis type stage structured predator–prey model. Applied Mathematics and Computation 244, 344–360.

Khan, J., Haq, E.U., Mehmood, T., Blouch, A., Rafi, M.A., Fateh, J., 2016a. Effect of temperature on the biology and predatory potential, of *Harmonia Dimidiata* (Fab.) (Coleoptera: Coccinellidae) feeding on *Myzus Persicae* (Sulzer) (Hemiptera: Aphididae) aphid. International Journal of Environment, Agriculture and Biotechnology 1, 342–349.

Khan, J., Haq, E.U., Rehman, A., 2015. Effect of temperature on the biology of *Harmonia dimidiate* Fab. (Coleoptera: Coccinellidae) reared on *Sciza-phus graminum* (Rond.) aphid. Journal of Biodiversity and Environmental Sciences 7, 42–49.

Khan, J., Haq, E.U., Saljoki, A.U.R., Rehman, A., 2016b. Effect of temperature on biological attributes and predatory potential of *Harmonia dimidi-ata* (Fab.)(Coleoptera: Coccinellidae) fed on *Rhopalosiphum padi* aphid. Journal of Entomology and Zoology Studies 4, 1016–1022.

Latham, D.R., Mills, N.J., 2010. Quantifying aphid predation: the mealy plum aphid hyalopterus pruni in california as a case study. Journal of Applied Ecology 47, 200–208.

Levins, R., 1966. The strategy of model building in population biology. American scientist 54, 421–431.

Lu, Y., Pawelek, K.A., Liu, S., 2017. A stage-structured predator-prey model with predation over juvenile prey. Applied Mathematics and Computation 297, 115–130.

Machekano, H., Mvumi, B.M., Nyamukondiwa, C., 2018. Loss of coevolved basal and plastic responses to temperature may underlie trophic level host-parasitoid interactions under global change. Biological Control 118, 44–54.

Mortoja, S.G., Panja, P., Mondal, S.K., 2018. Dynamics of a predator-prey model with stage-structure on both species and anti-predator behavior. Informatics in medicine unlocked 10, 50–57.

Mou, D.F., Lee, C.C., Smith, C., Chi, H., 2015. Using viable eggs to accurately determine the demographic and predation potential of harmonia dimidiata (c oleoptera: C occinellidae). Journal of applied entomology 139, 579–591.

Newman, J., 2005. Climate change and the fate of cereal aphids in Southern Britain. Global Change Biology 11, 940–944.

Newman, J., Gibson, D., Parsons, A., Thornley, J., 2003. How predictable are aphid population responses to elevated co2? Journal of Animal Ecology 72, 556–566.

Newman, J.A., 2004. Climate change and cereal aphids: the relative effects of increasing co2 and temperature on aphid population dynamics. Global Change Biology 10, 5–15.

Newman, J.A., 2006. Using the output from global circulation models to predict changes in the distribution and abundance of cereal aphids in Canada: a mechanistic modeling approach. Global Change Biology 12, 1634–1642.

Ng, J.C., Perry, K.L., 2004. Transmission of plant viruses by aphid vectors. Molecular Plant Pathology 5, 505–511.

Pintanel, P., Tejedo, M., Salinas-Ivanenko, S., Jervis, P., Merino-Viteri, A., 2021. Predators like it hot: Thermal mismatch in a predator–prey system across an elevational tropical gradient. Journal of Animal Ecology 90, 1985–1995.

Satar, S., Kersting, U., Uygun, N., 2005. Effect of temperature on development and fecundity of aphis gossypii glover (homoptera: Aphididae) on cucumber. Journal of Pest Science 78, 133–137.

Schmitz, O.J., Barton, B.T., 2014. Climate change effects on behavioral and physiological ecology of predator–prey interactions: implications for conservation biological control. Biological Control 75, 87–96.

Schulzweida, U., 2019. CDO user guide (Version 1.9.8). https://doi.org/10.5281/zenodo.3539275. doi:10.5281/zenodo.3539275. date accessed: October 31, 2019.

Sharma, P., Verma, S., Chandel, R., Shah, M., Gavkare, O., 2017. Functional response of *Harmonia dimidiata* (Fab.) to melon aphid, *Aphis gossypii*Glover under laboratory conditions. Phytoparasitica 45, 373–379.

Singh, R., Singh, K., 2015. Life history parameters of aphis gossypii glover (homoptera: Aphididae) reared on three vegetable crops. International Journal of Research Studies in Zoology 1, 1–9.

Skirvin, D., Perry, J., Harrington, R., 1997. The effect of climate change on an aphid–coccinellid interaction. Global change biology 3, 1–11.

Slosser, J., Parajulee, M., Hendrix, D., Henneberry, T., Pinchak, W., 2004. Cotton aphid (Homoptera: Aphididae) abundance in relation to cotton leaf sugars. Environmental Entomology 33, 690–699.

Sunday, J.M., Bates, A.E., Dulvy, N.K., 2012. Thermal tolerance and the global redistribution of animals. Nature Climate Change 2, 686–690.

Thornley, J.H., France, J., 2007. Mathematical models in agriculture: quantitative methods for the plant, animal and ecological sciences. Cabi.

Thornley, J.H., Newman, J.A., 2022. Climate sensitivity of the complex dynamics of the green spruce aphid - spruce plantation interactions: insight from a new mechanistic model. PLoS One, in press.

Tüzün, N., Stoks, R., 2018. Evolution of geographic variation in thermal performance curves in the face of climate change and implications for biotic interactions. Current opinion in insect science 29, 78–84.

Van Steenis, M., El-Khawass, K., 1995. Life history of aphis gossypii on cucumber: influence of temperature, host plant and parasitism. Entomologia Experimentalis et Applicata 76, 121–131.

Visser, M.E., Gienapp, P., 2019. Evolutionary and demographic consequences of phenological mismatches. Nature Ecology & Evolution 3, 879–885.

Wang, X., Zou, X., 2017. Modeling the fear effect in predator–prey interactions with adaptive avoidance of predators. Bulletin of mathematical biology 79, 1325–1359.

Xia, J., van der Werf, W., Rabbinge, R., 1999. Influence of temperature on bionomics of cotton aphid, aphis gossypii, on cotton. Entomologia Experimentalis et Applicata 90, 25–35.

Xia, J., Wang, J., Cui, J., Leffelaar, P., Rabbinge, R., Van Der Werf, W., 2018. Development of a stage-structured process-based predator–prey model to analyse biological control of cotton aphid, aphis gossypii, by the sevenspot ladybeetle, coccinella septempunctata, in cotton. Ecological complexity 33, 11–30.

Youngsteadt, E., Ernst, A.F., Dunn, R.R., Frank, S.D., 2017. Responses of arthropod populations to warming depend on latitude: evidence from urban heat islands. Global change biology 23, 1436–1447.

Yu, J., Chi, H., Chen, B.H., 2013. Comparison of the life tables and predation rates of *Harmonia dimidiata* (F.) (Coleoptera: Coccinellidae) fed on *Aphis gossypii* Glover (Hemiptera: Aphididae) at different temperatures. Biological Control 64, 1–9.

Zamani, A., Talebi, A., Fathipour, Y., Baniameri, V., 2006. Effect of temperature on biology and population growth parameters of *Aphis gossypii* Glover (Hom., Aphididae) on greenhouse cucumber. Journal of Applied Entomology 130, 453–460.

Zarghami, S., Mossadegh, M.S., Kocheili, F., Allahyari, H., Rasekh, A., 2016. Functional responses of *Nephus arcuatus* Kapur (Coleoptera: Coccinellidae), the most important predator of spherical mealybug *Nipaecoccus viridis* (Newstead). Psyche 2016, 1–9.

