## Supplements for "Predator-prey interactions in a warming world: the critical role of cold tolerance"

### SUPPLEMENTARY MATERIALS

#### Supplemental documents

**Supplement 1.** Life history of aphid and ladybird.

#### Supplemental Figures

Figure S1.Thermal performance curves (TPCs) for aphid's and ladybird's developmental rates and mortality rates under different thermal tolerance mismatch scenarios.

Figure S7. Heatmaps in parameter spaces for aphid and ladybird population abundance under different climate scenarios (*AL2: predators are more cold tolerant than the prey*, *A*-Class).

Figure S8. Heatmaps in parameter spaces for aphid and ladybird population abundance under different climate scenarios (*AL3: predators are more heat tolerant than the prey*, *A*-Class).

Figure S9. Heatmaps in parameter spaces for aphid and ladybird population abundance under different climate scenarios (*AL4: predators are more heat and more cold tolerant than the prey*, *A*-Class).

Figure S10. Heatmaps in parameter spaces for aphid and ladybird population abundance under different climate scenarios (*AL7: predators are both less cold and less heat tolerant than the prey*, *B*-Class).

Figure S11. Heatmaps in parameter spaces for aphid and ladybird population abundance under different climate scenarios (*AL8: predators are both more cold tolerant and more heat tolerant than the prey*, *B*-Class).

Figure S12. Heatmaps in parameter spaces for aphid and ladybird population abundance under different climate scenarios (*AL9: predators are less heat and more cold tolerant than the prey*, *B*-Class).

Figure S13. Spatial maps for the temperature metrics under current and future climates.

Figure S14. Heatmaps in parameter spaces for aphid and ladybird population abundance (*AAP* and *ALP*) under different thermal tolerance scenarios and different initial prey-predator ratio ($R_{initial}$).

#### Supplemental Tables

Table S1. Definitions of notations in aphid submodel.

Table S3. Definitions of notations in the ladybird submodel.

Table S2. Parameter values for the aphid submodel.

Table S4 Parameter values for the ladybird submodel (all nine ladybirds).

### SUPPLEMENT 1: LIFE HISTORY OF APHID AND LADYBIRD

#### *Aphis gossypii* life history

The life cycle of *Aphis gossypii* can include both parthenogenetic and sexual generations (Moran, 1992). In warmer environments, the aphids are primarily apterous (wingless) and exhibit an anholocyclic life cycle (summer cycle). After going through four instar nymph stages, asexual adult aphids give birth to live young. While some of the nymphs develop into alate (winged) adults and migrate to new host plants when the local density becomes too high, or the local plant quality deteriorates too much. However, in cooler environments, the aphids exhibit either a heteroecious or autoecious holocyclic life cycle (winter cycle; Moran, 1992). Since asexual reproduction occurs during most of the year, while sexual reproduction only occurs once during the autumn, most life table studies only focus on the anholocyclic life cycle (Ebert et al., 1997). The anholocyclic life cycle of *A. gossypii* is very short, the total development duration of the four instar nymphs is 3.5–8.2 d, and the longevity of apterous adults is 10.9–24.1 d (at a constant temperature of 25 °C; variation is due to the different host plants and aphid ecotypes) (Aldyhim and Khalil, 1993; Kocourek et al., 1994; Van Steenis and El-Khawass, 1995; Xia et al., 1999; Kersting et al., 1999; Satar et al., 2005; Zamani et al., 2006; Singh and Singh, 2015).

Many laboratory experiments show that aphid development rates of *A. gossypii* are temperature-dependent. The lower (5.8–10.47 °C) and upper (35–45 °C) developmental thresholds of this aphid depend on host plants and ecotypes (Kocourek et al., 1994; Ebert et al., 1997; Kersting et al., 1999; Xia et al., 1999; Satar et al., 2005; Zamani et al., 2006). Aldyhim and Khalil (1993) and Kersting et al. (1999) reported the optimum temperature range for growth and reproduction is 25–30 °C. Xia et al. (1999) reported that juvenile survival and fecundity rates were highest at 25 °C. Alate production is a complex interaction of several biotic and abiotic factors including plant quality, crowding, temperature, and photoperiod (De Barro, 1992; Liu, 1994). Ebert et al. (1997) conclude that "crowding is the driving force for alate production while other factors (e.g. nutrition, parentage, temperature) all modify the magnitude of the response." Hodgson et al. (2005) suggest that shorter photoperiods trigger alate production. Tegelaar and Leimar (2014) also investigated the influence of an alarm pheromone and ant attendance on alate production. How these factors combined to influence the alate production during the growing season is not fully understood.

#### *Harmonia dimidiata* life history

Unlike the aphid, the life cycle of the ladybird involves a complete metamorphosis. *Harmonia dimidiata* develops through the following stages: egg, four instar larvae, pupae, and adult. The development duration of each stage is much longer than *A. gossypii*. For *H. dimidiata* that fed on *A. gossypii* at a constant temperature of 25 °C, the development durations of its egg, four instar larvae, pupae, and adult were: 3.41–4.00, 2.0–2.10, 1.37–1.60,

1.70–1.90, 5.50–5.75, 3.60–5.07 and 55.80–67.73 days, respectively (Yu et al., 2013; Mou et al., 2015).

Previous laboratory experiments have demonstrated that "temperature has a profound effect on the developmental durations, survival, fecundity, and predatory potential of *H. dimidiata*." (Khan et al., 2015, p.42). The most favorable temperature for culturing *H. dimidiata* was 20–25°C (Kuznetsov and Pang, 2002; Agarwala et al., 2009; Khan et al., 2015). In the laboratory experiments at five constant temperatures (15, 20, 25, and 30°C or 16, 20, 24, 28, and 32°C), maximum developmental durations of different stages were observed at low temperature, and minimum developmental durations were observed at high temperature (Khan et al., 2016a,b). The female ladybird does not produce eggs when the temperature is above 30°C (Yu et al., 2013; Khan et al., 2015). The maximal daily fecundity of females increased with temperature, while the maximum lifetime fecundity was higher at lower temperatures (Yu et al., 2013). As the dominant predator of many aphid species, both larvae and adults of *H. dimidiata* are highly voracious, the adult beetles and the fourth-instar were the most voracious stages. They can consume more than 200 aphids per day (Yu et al., 2013; Mou et al., 2015). The transformation rate ($Q_p$), the number of aphids that the predator needs to eat to produce one offspring, is estimated to be 74.3–131.8 aphids/female. Consumption rates are maximal at 20 °C (Yu et al., 2013; but see Mou et al., 2015).

### REFERENCES


Agarwala, B. K., Singh, T. K., Lokeshwari, R. K., and Sharmila, M. (2009). Functional response and reproductive attributes of the aphidophagous ladybird beetle, *Harmonia dimidiata* (Fabricius) in oak trees of sericultural importance. *Journal of Asia-Pacific Entomology*, 12(3):179–182.

Aldyhim, Y. and Khalil, A. (1993). Influence of temperature and daylength on population development of *Aphis gossypii* on *Cucurbita pepo*. *Entomologia Experimentalis et Applicata*, 67(2):167–172.

De Barro, P. (1992). The role of temperature, photoperiod, crowding and plant quality on the production of alate viviparous females of the bird cherry-oat aphid, *Rhopalosiphum padi*. *Entomologia Experimentalis et Applicata*, 65(3):205–214.

Ebert, T., Cartwright, B., et al. (1997). Biology and ecology of *Aphis gossypii* Glover (Homoptera: Aphididae). *Southwestern Entomologist*, 22(1):116–153.

Hodgson, E., Venette, R., Abrahamson, M., and Ragsdale, D. (2005). Alate production of soybean aphid (Homoptera: Aphididae) in Minnesota. *Environmental Entomology*, 34(6):1456–1463.

Kersting, U., Satar, S., and Uygun, N. (1999). Effect of temperature on development rate and fecundity of apterous *Aphis gossypii* Glover (Hom., Aphididae) reared on *Gossypium hirsutum* L. *Journal of Applied Entomology*, 123(1):23–27.

Khan, J., Haq, E. U., Mehmood, T., Blouch, A., Rafi, M. A., and Fateh, J. (2016a). Effect of temperature on the biology and predatory potential, of *Harmonia Dimidiata* (Fab.) (Coleoptera: Coccinellidae) feeding on

*Myzus Persicae* (Sulzer) (Hemiptera: Aphididae) aphid. *International Journal of Environment, Agriculture and Biotechnology*, 1(3):342–349.

Khan, J., Haq, E. U., and Rehman, A. (2015). Effect of temperature on the biology of *Harmonia dimidiate* Fab. (Coleoptera: Coccinellidae) reared on *Scizaphus graminum* (Rond.) aphid. *Journal of Biodiversity and Environmental Sciences*, 7:42–49.

Khan, J., Haq, E. U., Saljoki, A. U. R., and Rehman, A. (2016b). Effect of temperature on biological attributes and predatory potential of *Harmonia dimidiata* (Fab.)(Coleoptera: Coccinellidae) fed on *Rhopalosiphum padi* aphid. *Journal of Entomology and Zoology Studies*, 4:1016–1022.

Kocourek, F., Havelka, J., Berankova, J., and Jaroŝik, V. (1994). Effect of temperature on development rate and intrinsic rate of increase of *Aphis gossypii* reared on greenhouse cucumbers. *Entomologia Experimentalis et Applicata*, 71(1):59–64.

Kuznetsov, V. and Pang, H. (2002). Employment of Chinese Coccinellidae in biological control of aphids in greenhouse in Primorye. *Far Eastern Entomologist*, pages 1–5.

Liu, S.-S. (1994). Production of alatae in response to low temperature in aphids: a trait of seasonal adaptation. In *Insect life-cycle polymorphism*, pages 245–261. Springer.

Moran, N. A. (1992). The evolution of aphid life cycles. *Annual Review of Entomology*, 37(1):321–348.

Mou, D.-F., Lee, C.-C., Smith, C., and Chi, H. (2015). Using viable eggs to accurately determine the demographic and predation potential of *Harmonia dimidiata* (Coleoptera: Coccinellidae). *Journal of Applied Entomology*, 139(8):579–591.

Satar, S., Kersting, U., and Uygun, N. (2005). Effect of temperature on development and fecundity of *Aphis gossypii* glover (Homoptera: Aphididae) on cucumber. *Journal of Pest Science*, 78(3):133–137.

Singh, R. and Singh, K. (2015). Life history parameters of *Aphis gossypii* Glover (Homoptera: Aphididae) reared on three vegetable crops. *International Journal of Research Studies in Zoology*, 1(1):1–9.

Tegelaar, K. and Leimar, O. (2014). Alate production in an aphid in relation to ant tending and alarm pheromone. *Ecological Entomology*, 39(5):664–666.

Van Steenis, M. and El-Khawass, K. (1995). Life history of *Aphis gossypii* on cucumber: influence of temperature, host plant and parasitism. *Entomologia Experimentalis et Applicata*, 76(2):121–131.

Xia, J., van, der Werf, W., and Rabbinge, R. (1999). Influence of temperature on bionomics of cotton aphid, *Aphis gossypii*, on cotton. *Entomologia Experimentalis et Applicata*, 90(1):25–35.

Yu, J., Chi, H., and Chen, B.-H. (2013). Comparison of the life tables and predation rates of *Harmonia dimidiata* (F.) (Coleoptera: Coccinellidae) fed on *Aphis gossypii* Glover (Hemiptera: Aphididae) at different temperatures. *Biological Control*, 64(1):1–9.

Zamani, A., Talebi, A., Fathipour, Y., and Baniameri, V. (2006). Effect of temperature on biology and population growth parameters of *Aphis gossypii* Glover (Hom., Aphididae) on greenhouse cucumber. *Journal of Applied Entomology*, 130(8):453–460.

### SUPPLEMENTAL FIGURES

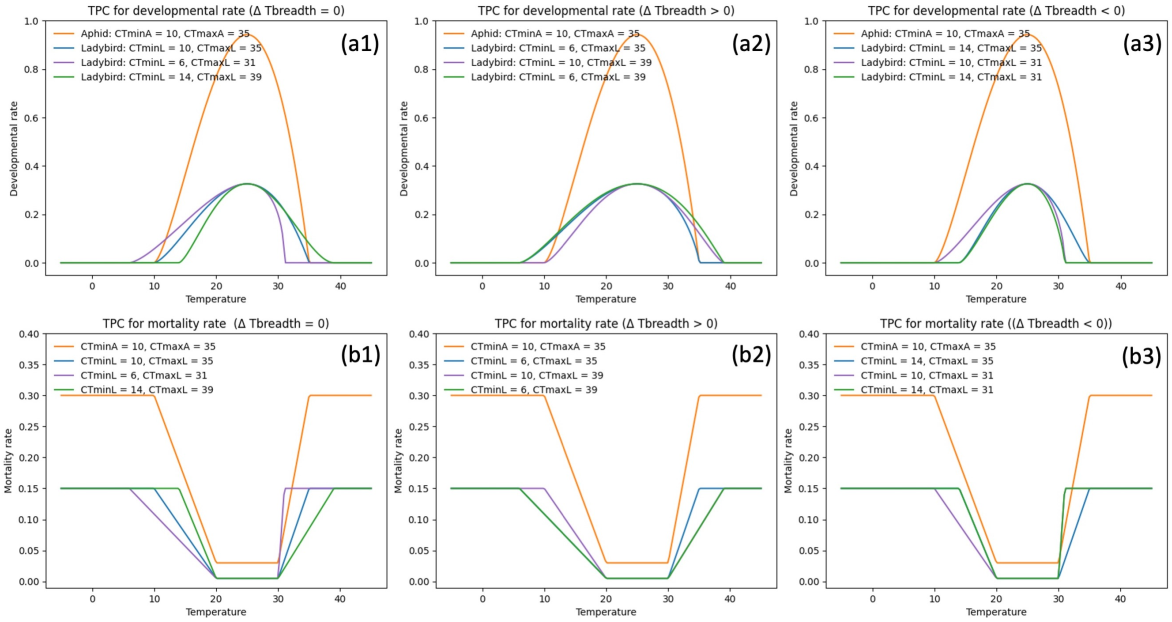


**Figure S1.** Thermal performance curves (TPCs) for aphid's and ladybird's developmental rates and mortality rates under different thermal tolerance mismatch scenarios. (a1)-(a3) represent the TPCs for developmental rates, and (b1)-(b3) represent the TPCs for the mortality rates. The orange curves in these plots represent the TPCs for the aphid, the rest curves represent TPCs for different ladybirds which have different thermal tolerances. The three ladybirds in (a1) and (b1) have the same thermal breadth as the aphid ($\Delta T_{breadth} = 0$), the three ladybirds in (a2) and (b2) have a wider thermal breadth ($\Delta T_{breadth} > 0$), while those in (a3) and (b3) have a narrower thermal breadth ($\Delta T_{breadth} < 0$).

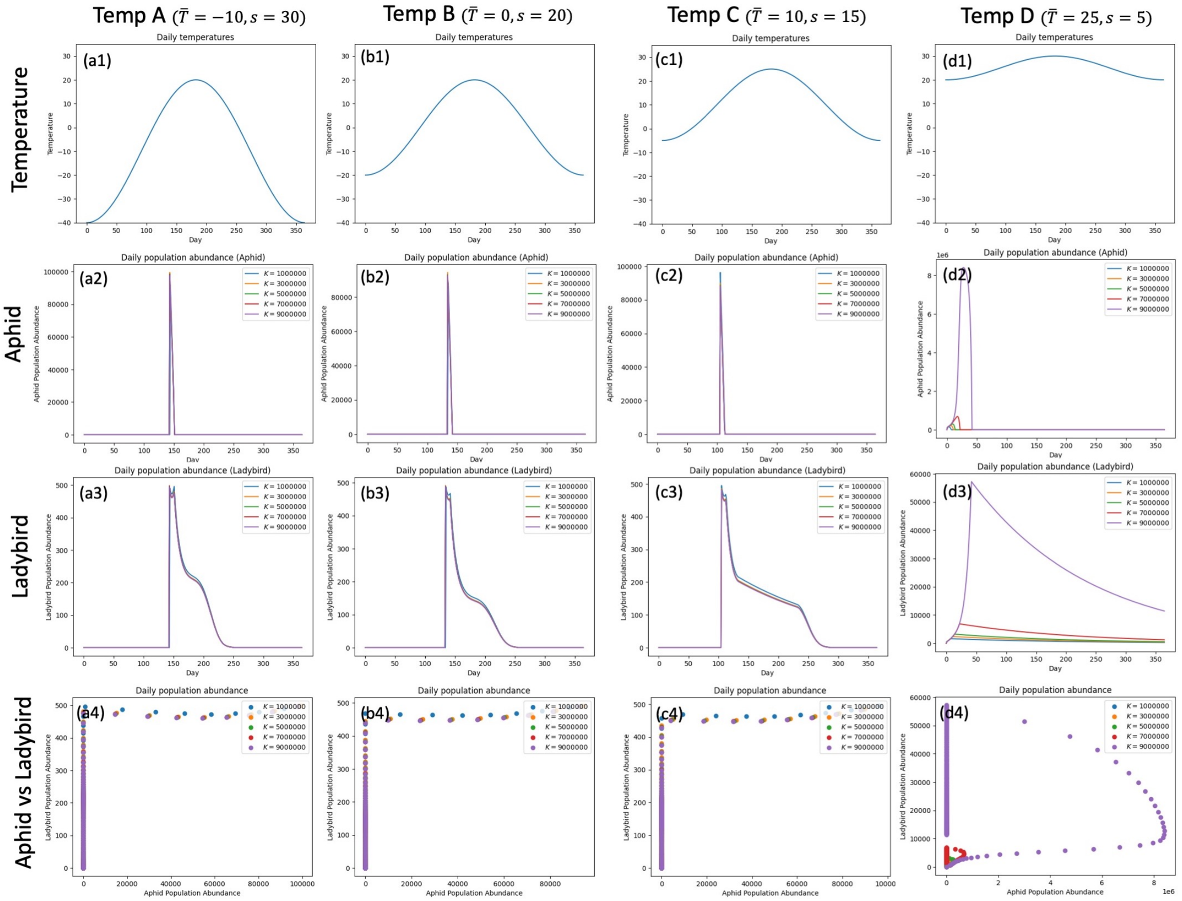


**Figure S2.** Sensitivity analysis for carrying capacity of aphid ($K$). The range of $K$ is set from $1 \times 10^6$ to $9 \times 10^6$. (a1)–(d1) represent the different temperature curves under four different climate conditions (four combinations of yearly mean temperature $\bar{T}$ and seasonality $s$). The subplots (a2)–(d4) summarize the simulations of aphid and ladybird daily population abundance with different $K$ under four different climate conditions. These plots indicate that altering the $K$ won't strongly affect the population dynamics of the aphid and ladybird, except when both species live in warmer less seasonal climate.

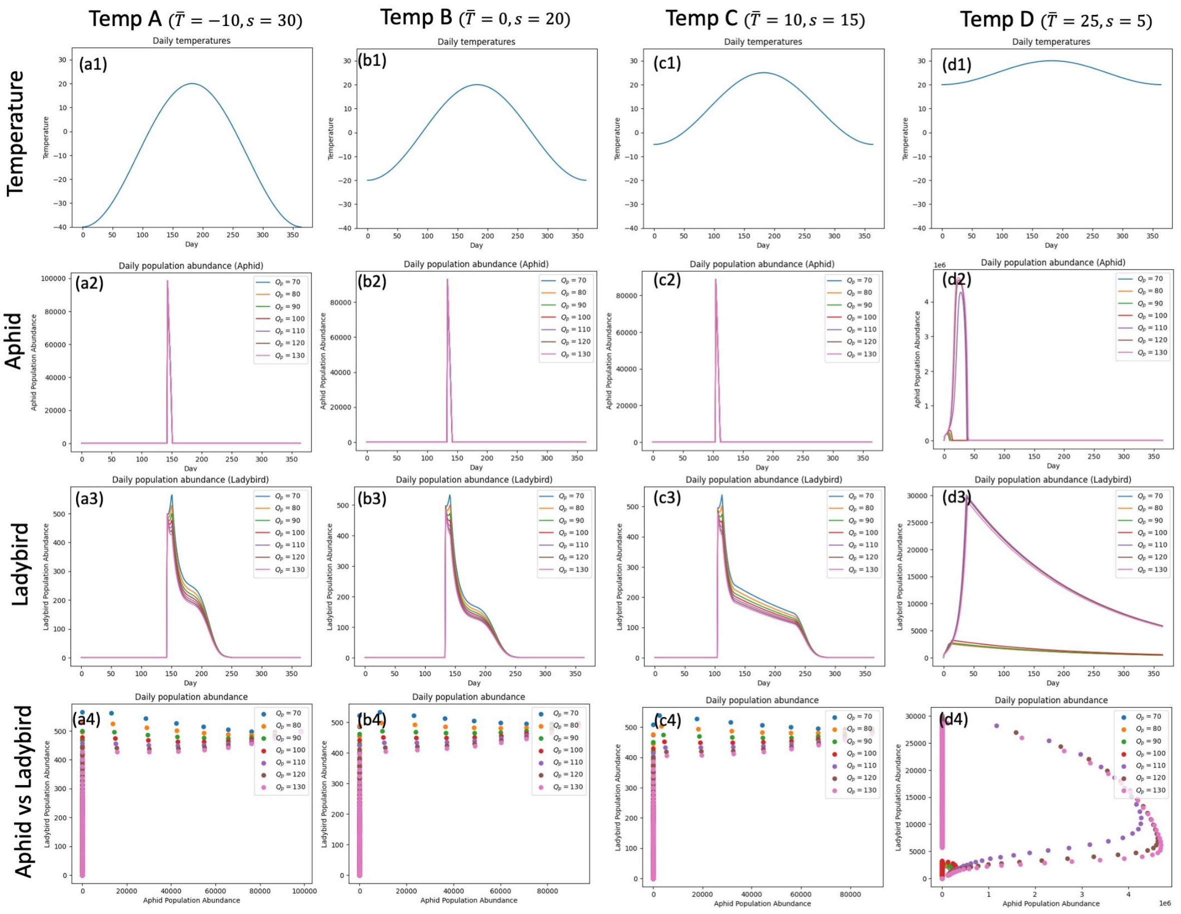


**Figure S3.** Sensitivity analysis for the transformation rate of ladybird ($Q_p$). The range of $Q_p$ is set from 70 to 130. (a1)–(d1) represent the different temperature curves under four different climate conditions (four combinations of yearly mean temperature $\bar{T}$ and seasonality $s$). The subplots (a2)–(d4) summarize the simulations of aphid and ladybird daily population abundance with different $Q_p$ under four different climate conditions. These plots indicate that altering the $Q_p$ won't strongly affect the population dynamics of the aphid and ladybird, except when both species live in warmer less seasonal climate.

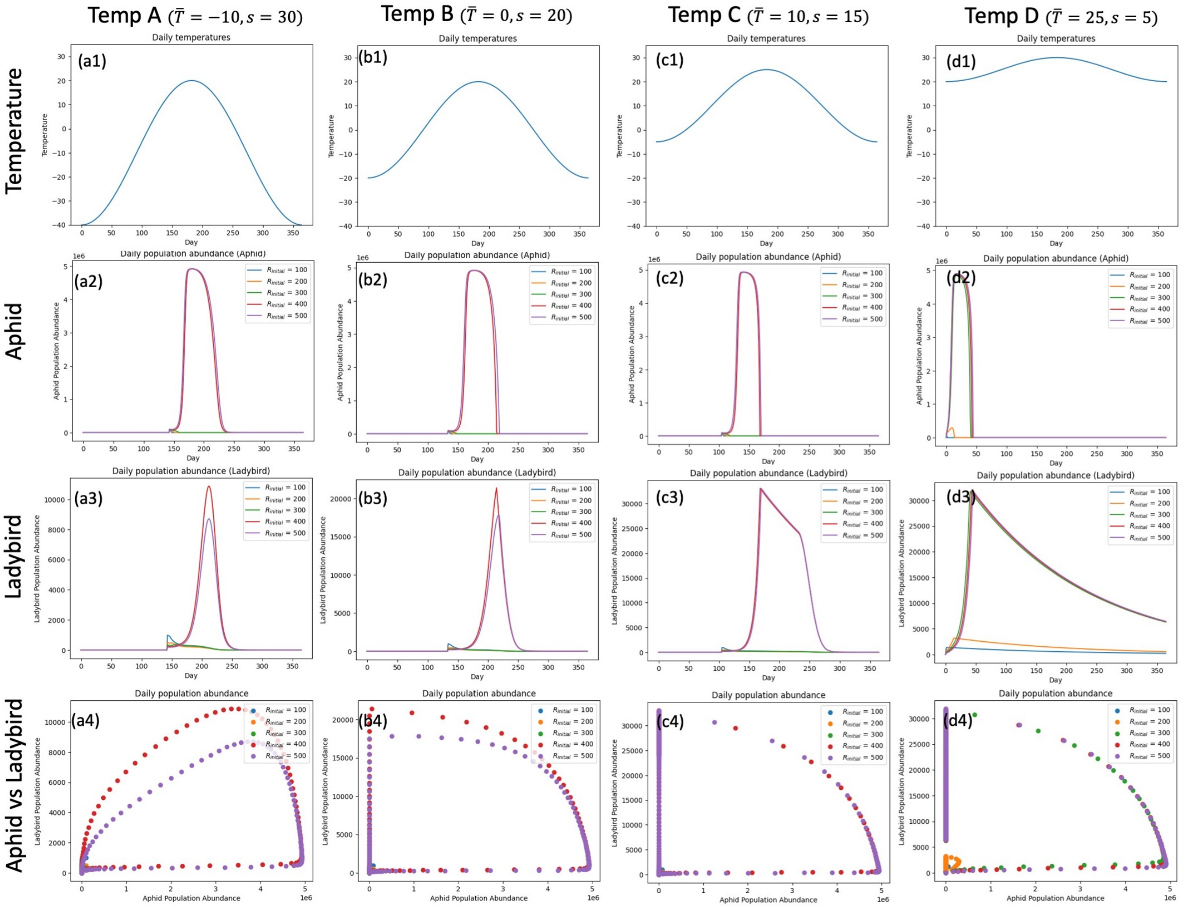


**Figure S4.** Sensitivity analysis for the ratio of aphid's initial value on ladybird's $R_{initial}$. The range of $R_{initial}$ is set from 100 to 500, the initial value for aphid is set to be a constant value ($1 \times 10^5$), while the initial values for ladybird are from 200 to 500, respectively. The subplots (a2)–(d4) summarize the simulations of aphid and ladybird daily population abundance with different $R_{initial}$ under five different climate conditions. These plots indicate that altering the $R_{initial}$ will strongly affect the population dynamics of the aphid and ladybird.

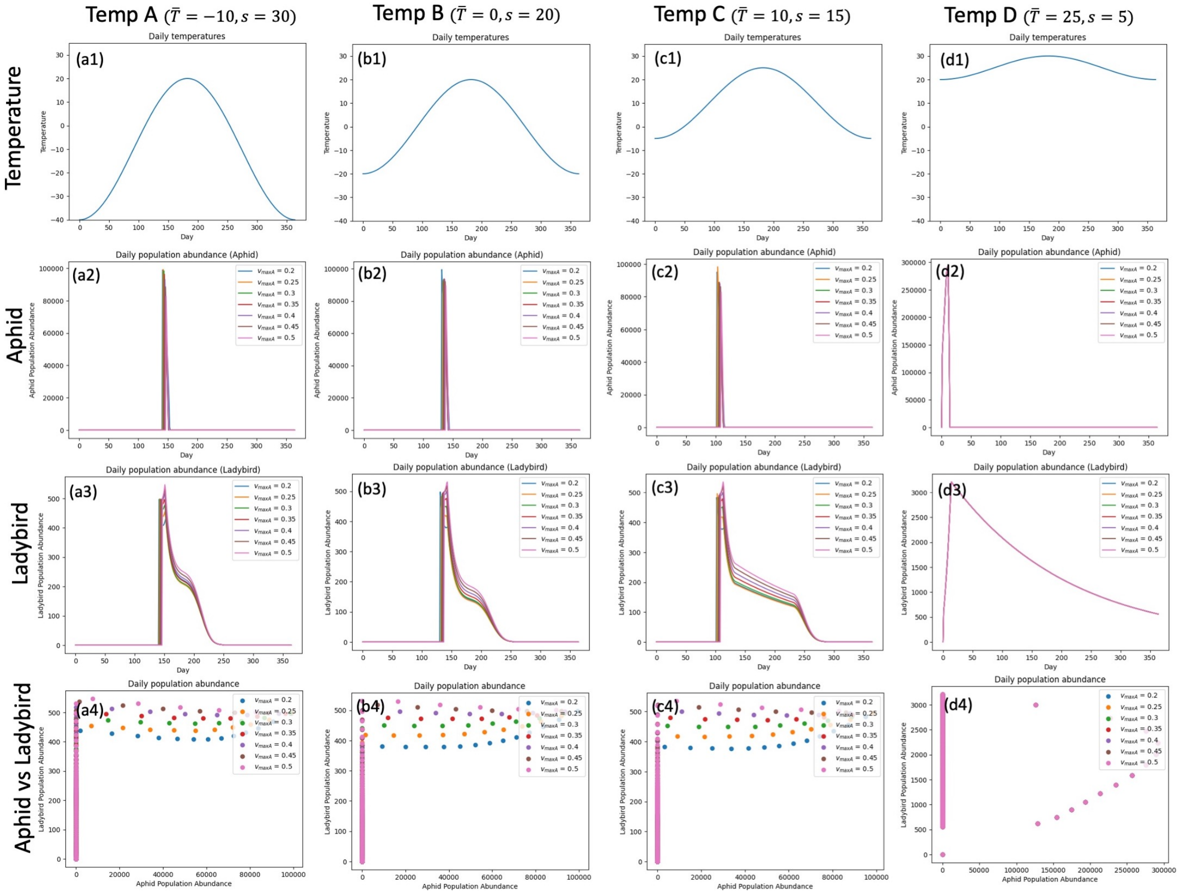


**Figure S5.** Sensitivity analysis for maximum mortality rate of aphid ($\upsilon_{maxA}$). The range of $\upsilon_{maxA}$ is set from 0.2 to 0.5. The subplots (a2)–(d4) summarize the simulations of aphid and ladybird daily population abundance with different $\upsilon_{maxA}$ under five different climate conditions. These plots indicate that altering the $\upsilon_{maxA}$ won't strongly affect the population dynamics of the aphid and ladybird.

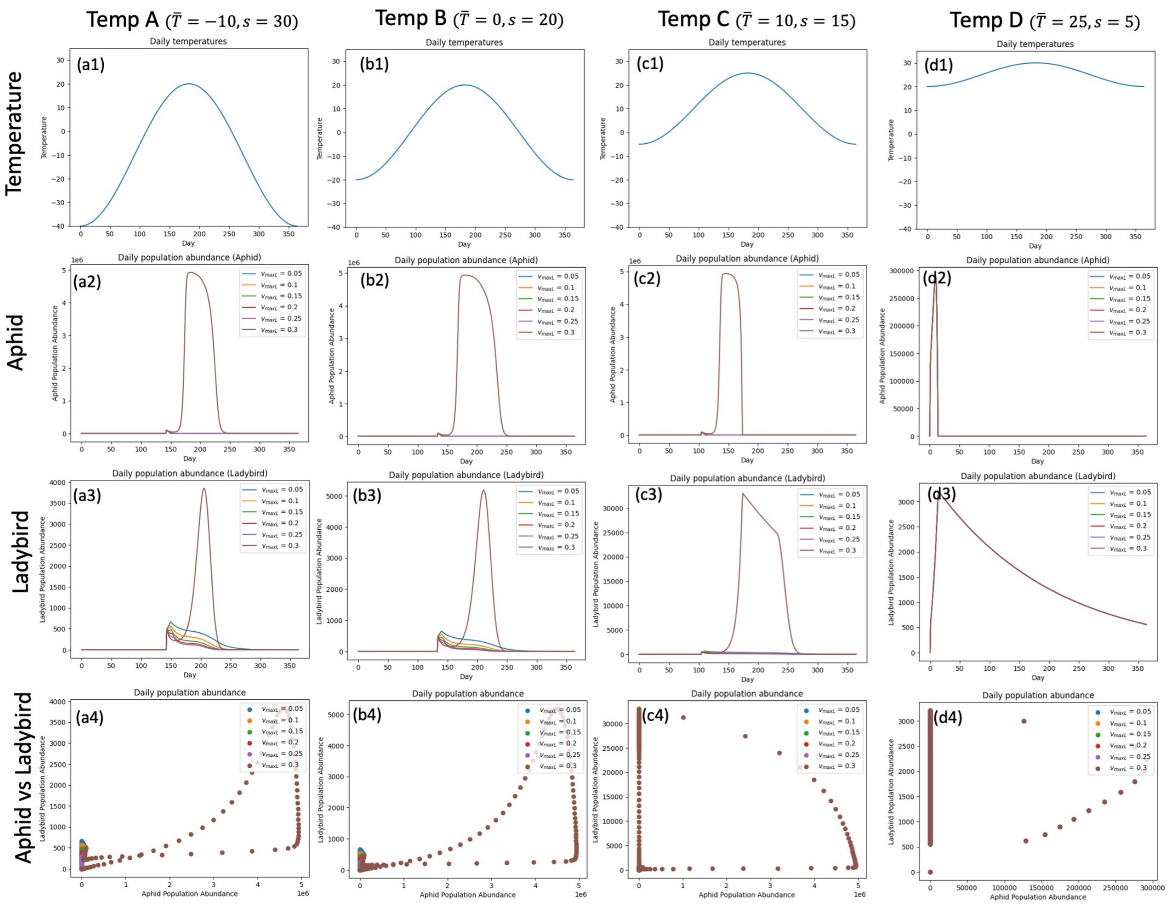


**Figure S6.** Sensitivity analysis for maximum mortality rate of aphid ($\upsilon_{maxL}$). The range of $\upsilon_{maxL}$ is set from 0.2 to 0.5. The subplots (a2)–(d4) summarize the simulations of aphid and ladybird daily population abundance with different $\upsilon_{maxL}$ under five different climate conditions. These plots indicate that altering the $\upsilon_{maxL}$ won't strongly affect the population dynamics of the aphid and ladybird. Note as well that the results are more sensitive to $\upsilon_{maxL}$ than $\upsilon_{maxA}$ (Figure S5) .

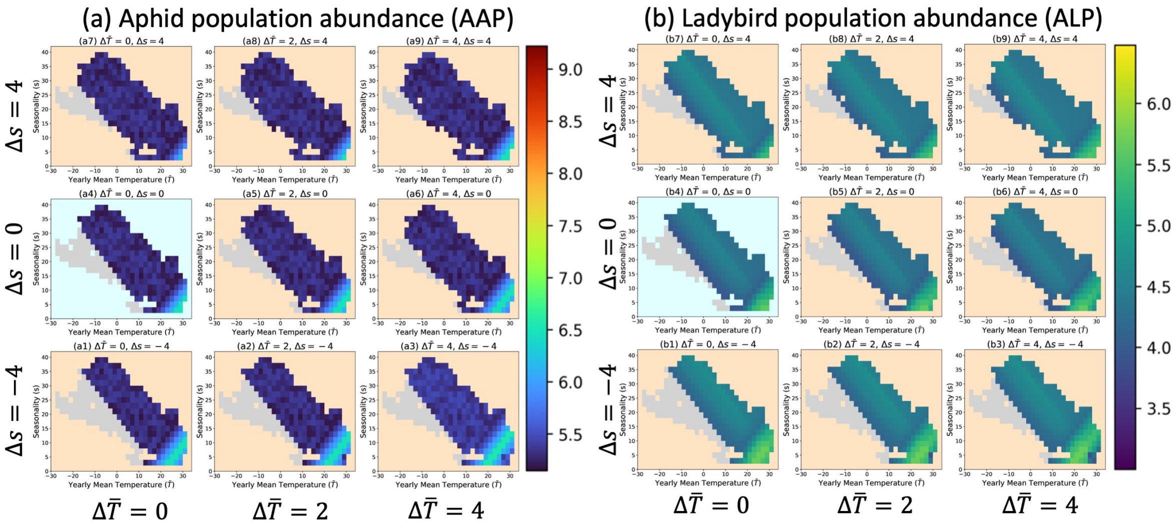


**Figure S7.** Heatmaps in parameter spaces for aphid and ladybird population abundance under different climate scenarios (*AL2: predators are more cold tolerant than the prey*, *A*-Class). Colored region represents the global combinations for mean temperature and seasonality. The left nine heatmaps (a) represent the patterns for aphid abundance. The right nine heatmaps (b) represent the patterns for ladybird abundance. The points with light grey color indicate that climates are unsuitable for the prey or predator. The heatmap with light blue background represents the abundance pattern under current climate, the rest heatmaps with light orange backgrounds represent the abundance patterns under different future climates.

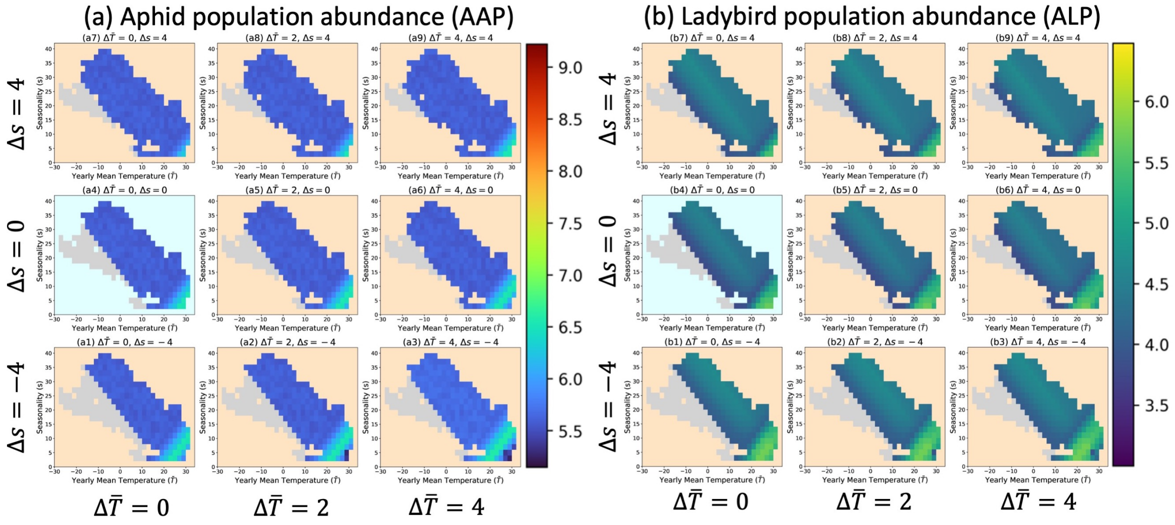


**Figure S8.** Heatmaps in parameter spaces for aphid and ladybird population abundance under different climate scenarios (*AL3: predators are more heat tolerant than the prey*, *A*-Class). Colored region represents the global combinations for mean temperature and seasonality. The left nine heatmaps (a) represent the patterns for aphid abundance. The right nine heatmaps (b) represent the patterns for ladybird abundance. The points with light grey color indicate that climates are unsuitable for the prey or predator. The heatmap with light blue background represents the abundance pattern under current climate, the rest heatmaps with light orange backgrounds represent the abundance patterns under different future climates.

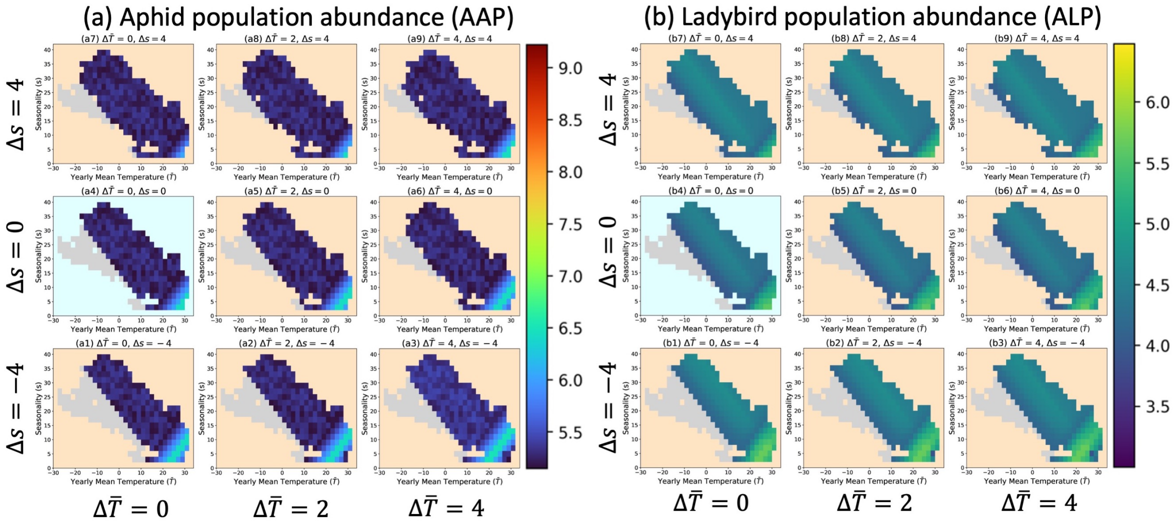


**Figure S9.** Heatmaps in parameter spaces for aphid and ladybird population abundance under different climate scenarios (*AL4: predators are more heat and more cold tolerant than the prey*, *A*-Class). Colored region represents the global combinations for mean temperature and seasonality. The left nine heatmaps (a) represent the patterns for aphid abundance. The right nine heatmaps (b) represent the patterns for ladybird abundance. The points with light grey color indicate that climates are unsuitable for the prey or predator. The heatmap with light blue background represents the abundance pattern under current climate, the rest heatmaps with light orange backgrounds represent the abundance patterns under different future climates.

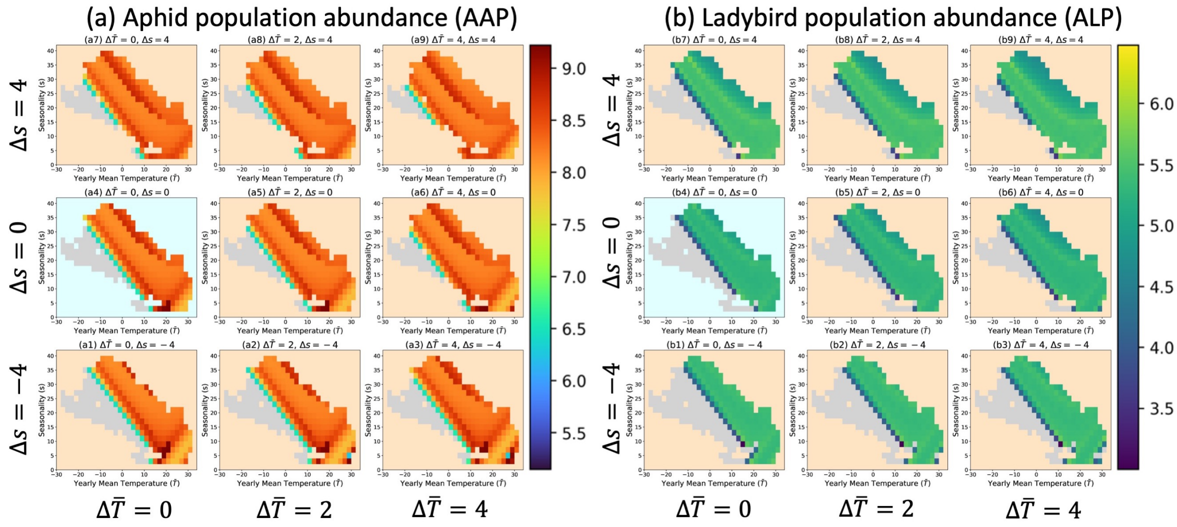


**Figure S10.** Heatmaps in parameter spaces for aphid and ladybird population abundance under different climate scenarios (*AL7: predators are both less cold and less heat tolerant than the prey*, *B*-Class). Colored region represents the global combinations for mean temperature and seasonality. The left nine heatmaps (a) represent the patterns for aphid abundance. The right nine heatmaps (b) represent the patterns for ladybird abundance. The points with light grey color indicate that climates are unsuitable for the prey or predator. The heatmap with light blue background represents the abundance pattern under current climate, the rest heatmaps with light orange backgrounds represent the abundance patterns under different future climates.

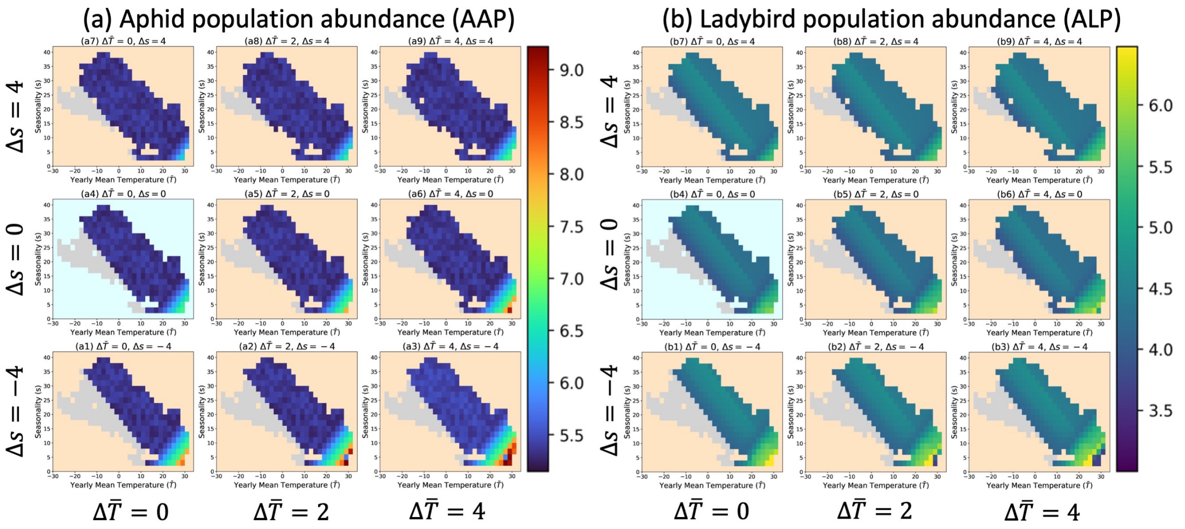


**Figure S11.** Heatmaps in parameter spaces for aphid and ladybird population abundance under different climate scenarios (*AL8: predators are both more cold tolerant and more heat tolerant than the prey*, *C*-Class). Colored region represents the global combinations for mean temperature and seasonality. The left nine heatmaps (a) represent the patterns for aphid abundance. The right nine heatmaps (b) represent the patterns for ladybird abundance. The points with light grey color indicate that climates are unsuitable for the prey or predator. The heatmap with light blue background represents the abundance pattern under current climate, the rest heatmaps with light orange backgrounds represent the abundance patterns under different future climates.

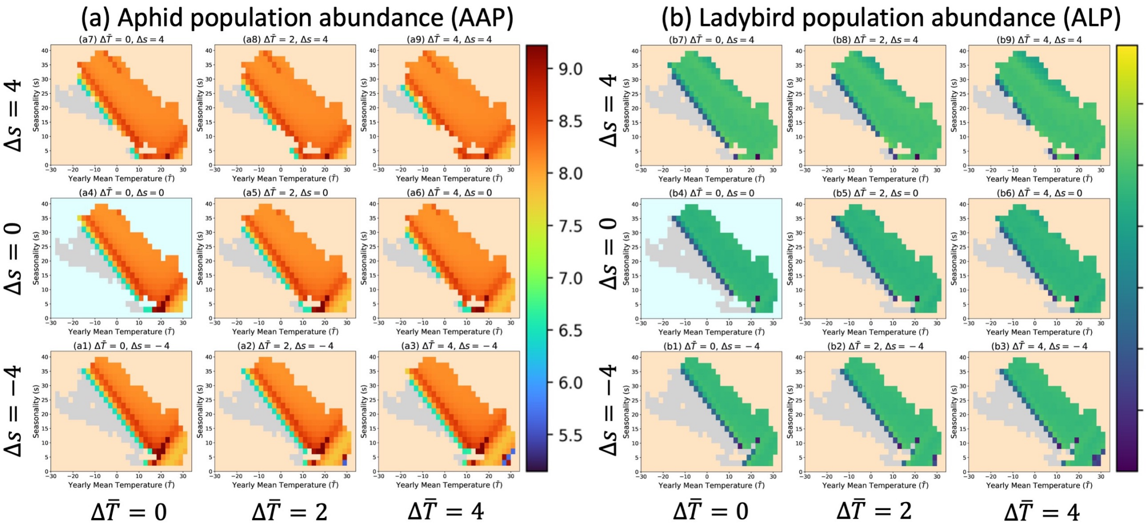


**Figure S12.** Heatmaps in parameter spaces for aphid and ladybird population abundance under different climate scenarios (*AL9: predators are less heat and more cold tolerant than the prey*, *B*-Class). Colored region represents the global combinations for mean temperature and seasonality. The left nine heatmaps (a) represent the patterns for aphid abundance. The right nine heatmaps (b) represent the patterns for ladybird abundance. The points with light grey color indicate that climates are unsuitable for the prey or predator. The heatmap with light blue background represents the abundance pattern under current climate, the rest heatmaps with light orange backgrounds represent the abundance patterns under different future climates.

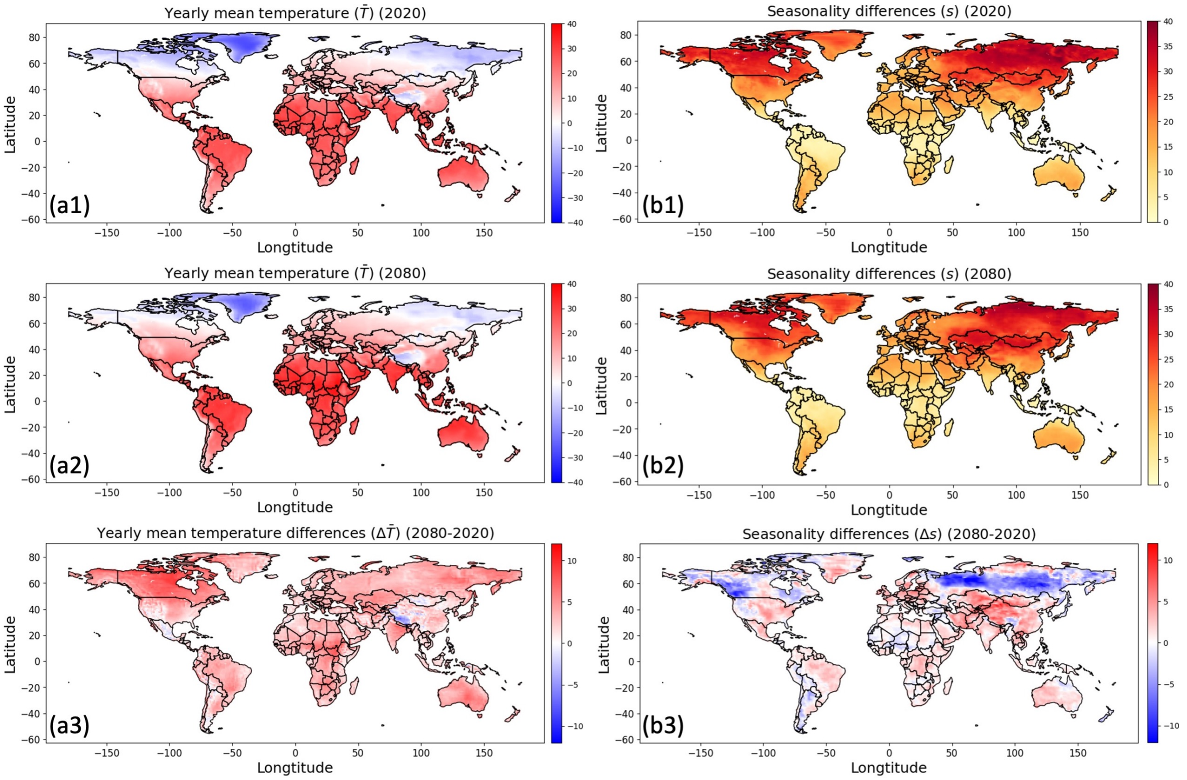


**Figure S13.** Spatial maps for the temperature metrics under current and future climates. (a1) and (a2) represent the spatial maps for the yearly mean temperatures ($\bar{T}$) in 2020 and 2080, and (a3) represents the difference between them ($\Delta\bar{T} = \bar{T}_{2080} - \bar{T}_{2020}$). (b1) and (b2) represent the spatial maps for the seasonality ($s$) in 2020 and 2080, and (b3) represents their difference ($\Delta s = s_{2080} - s_{2020}$). Please see section 2.3 for the details about how we get the yearly mean temperature.

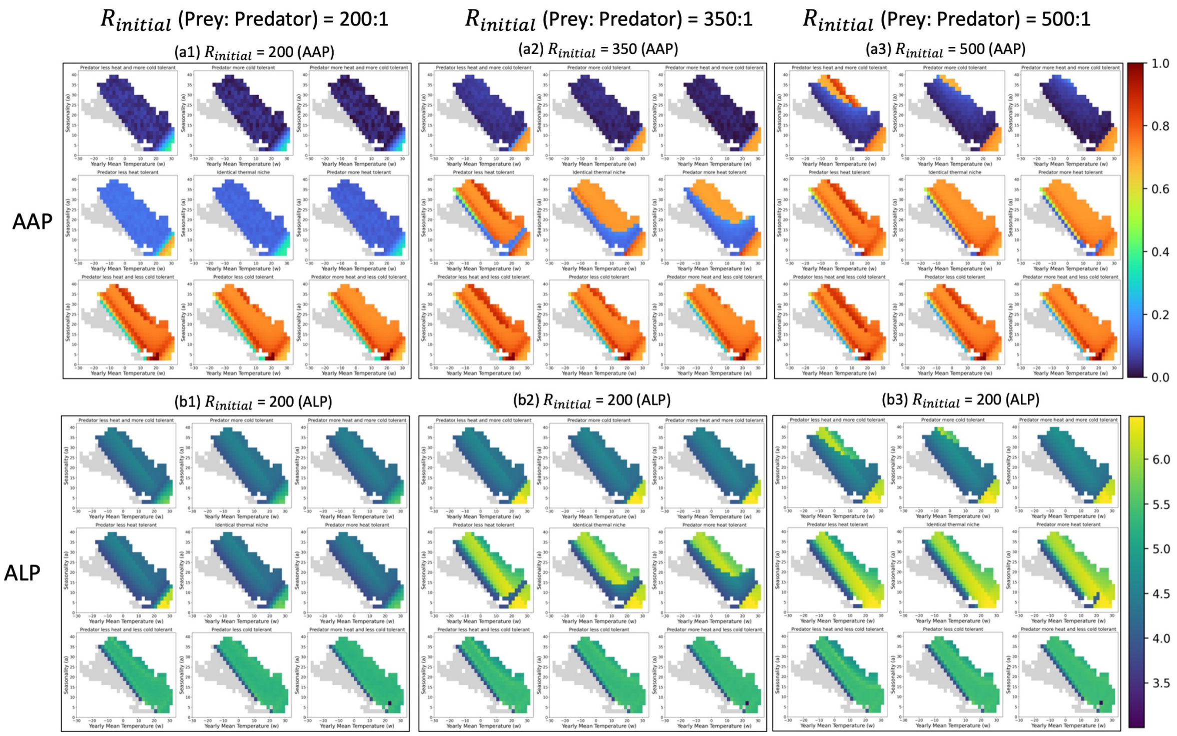


**Figure S14.** Heatmaps in parameter spaces for aphid and ladybird population abundance (*AAP* and *ALP*) under different thermal tolerance scenarios and different initial prey-predator ratio ($R_{initial}$). The 9 panels in each plot (e.g., (a1)) represent the results under different thermal tolerance scenarios. (a1)-(a3) represent the *AAP* heatmaps when the initial prey-predator ratios are 200, 350, and 500. (b1)-(b3) represent the *ALP* heatmaps under three different $R_{initial}$.

### SUPPLEMENTAL TABLES

**Table S1.** Definitions of notations in aphid submodel.

| Notation | Definition |
|---|---|
| $f_{ap}(T)$ | Fecundity rate of apterous adult, denotes the total number of nymphs produced by per apterous adult per day. |
| $\varphi(T)$ | Development rate of the each nymph, which is the fraction of time spent in each stage. |
| $\mu_{ny}(T)$, $\mu_{ap}(T)$ | Mortality rate of aphids (instar numphs and apterous adults), denotes per capita daily intrinsic mortality. |
| $P_i$ | Predation rate on aphids (instars and apterous adults), denotes the number of stage specific aphids captured by the ladybird population per day. $i$ represent different life stage of the aphid. |
| $A_{den}$ | Total aphid population density (The initial value for the model is assumed as $1 \times 10^5$ aphid individuals per $100m^2$). |
| $A_i$ | Stage specific aphid population density. |
| $a_i$ | Fraction of stage specific aphid in the total population, denotes its relative abundance in the total aphid population. |
| $P$ | Total number of aphid consumed by the ladybird population per day. |
| $\sigma_i$ | Stage specific predation pressure for per capita aphid. |
| $K$ | Carrying capacity of aphid ($5 \times 10^6$ aphid individuals per 100 $m^2$). |
| $CT_{min}$,$CT_{max}$ | The critical thermal minimum and maximum are temperature thresholds beyond which fecundity rate and developmental rate are minimized to be 0, or mortality rate reaches to maximum. |
| $CT_{opt}$, $CT_{opt1}$, $CT_{opt2}$ | $CT_{opt}$ is the optimal temperature for the development of aphid. The developmental rate and fecundity rate reach to their maximum at $CT_{opt}$. $CT_{opt1}$ and $CT_{opt2}$ are the thresholds of optimal temperature range for aphid. |
| $m$,$q_1$, $q_2$ | Shape parameters used in Eq. 1 for aphid's fecundity rate and developmental rate. |
| $a_1$, $a_2$, $b_1$, $b_2$ | Shape parameters used in Eq. 3 for aphid's mortality rate. |

**Table S2.** Parameter values for the aphid submodel.

| Fecundity | Development | Mortality |
|---|---|---|
| $m_{fec} = 4.848$ | $m_{dev} = 0.9421$ | $v_{min} = 0.03$ |
| $q_{1,fec} = 1.5$ | $q_{1,dev} = 1.5$ | $v_{max} = 0.3$ |
| $q_{2,fec} = 1$ | $q_{2,dev} = 1$ | $k_1 = -0.027$ |
| $CT_{min} = 10$ | $CT_{min} = 10$ | $b_1 = 0.57$ |
| $CT_{max} = 35$ | $CT_{max} = 35$ | $k_2 = 0.054$ |
| | | $b_2 = -1.59$ |
| | | $CT_{opt1} = 20$ |
| | | $CT_{opt2} = 30$ |

**Table S3.** Definitions of notations in the ladybird submodel.

| Notation | Definition |
|---|---|
| $f_H(T)$ | Fecundity rate for female ladybird, denotes the total number of eggs produced by per female adult per day. |
| $\beta_j(A_{den}, T)$ | Predation rate of stage-specific ladybird, denotes the number of aphids eaten by per ladybird per day. |
| $\delta_j(T)$ | Developmental rate for egg and pupa of ladybird. Depends on the ambient air temperature. |
| $\delta_j(A_{den}, T)$ | Developmental rate for different larvae stage of ladybird. Depends on the ambient air temperature and prey density. |
| $\gamma_j(T)$ | Mortality rate of each stage of ladybird, denotes per capita daily intrinsic mortality. |
| $G_\beta(A^*_{den}, T)$ | Temperature-dependent predation rate at saturation. |
| $g(A^*_{den}, T)$ | An index ranging from 0 to 1 that scales the predation rate function depending in temperature. |
| $H_{den}$ | Total population density of ladybird that can consume aphid. The initial value is assumed as 200 ladybird beetles per $100m^2$. |
| $H_j$ | Population density of the vulnerable stages of ladybird. |
| $CT_{min}$,$CT_{max}$ | The critical thermal minimum and maximum are temperature thresholds beyond which fecundity rate and developmental rate are minimized to be 0, or mortality rate reaches to maximum. |
| $CT_{opt}$, $CT_{opt1}$, $CT_{opt2}$ | $CT_{opt}$ is the optimal temperature for the development of the ladybird. The developmental rate and fecundity rate reach to their maximum at $CT_{opt}$. $CT_{opt1}$ and $CT_{opt2}$ are the thresholds of optimal temperature range for the ladybird. |
| $m$,$q_1$, $q_2$ | Shape parameters used in Eq. 1 for ladybird's fecundity rate and developmental rate. |
| $a_1$, $a_2$, $b_1$, $b_2$ | Shape parameters used in Eq. 3 for ladybird's mortality rate. |
| $a_j$ | Searching time for the ladybird to encounter an aphid. |
| $h_j$ | Handling time for the ladybird to process an aphid. |
| $Q_p$ | Transformation rate, denotes the mean number of aphids a female ladybird beetle needs to consume to produce a single egg (100 aphid individuals per ladybird. |
| $\theta$ | A proportion of adult ladybird that are females. |

**Table S4.** Parameter values for the ladybird submodel (all nine ladybirds).

| Class | Ladybird1 | Ladybird2 | Ladybird3 | Ladybird4 | Ladybird5 | Ladybird6 | Ladybird7 | Ladybird8 | Ladybird9 |
|---|---|---|---|---|---|---|---|---|---|
| Thermal tolerance | $CT_{min} = 10$ | $CT_{min} = 6$ | $CT_{min} = 10$ | $CT_{min} = 6$ | $CT_{min} = 14$ | $CT_{min} = 10$ | $CT_{min} = 14$ | $CT_{min} = 6$ | $CT_{min} = 14$ |
| | $CT_{max} = 35$ | $CT_{max} = 35$ | $CT_{max} = 39$ | $CT_{max} = 39$ | $CT_{max} = 35$ | $CT_{max} = 31$ | $CT_{max} = 31$ | $CT_{max} = 31$ | $CT_{max} = 39$ |
| Development | $T^* = 25$, $m = 0.326$, $q_1 = 1.5$ | | | | | | | | |
| | $q_2$ = 1 | $q_2$ = 0.7895 | $q_2$ = 1.4 | $q_2$ = 1.105 | $q_2$ = 1.364 | $q_2$ = 0.6 | $q_2$ = 0.8182 | $q_2$ = 0.4737 | $q_2$ = 1.909 |
| Mortality | $CT_{opt1} = 20$, $CT_{opt2} = 30$, $v_{max} = 0.15$, $v_{min} = 0.005$ | | | | | | | | |
| | $k_1$ = -0.0145 | $k_1$ = -0.01036 | $k_1$ = -0.0145 | $k_1$ = -0.1036 | $k_1$ = -0.02417 | $k_1$ = -0.0145 | $k_1$ = -0.02417 | $k_1$ = -0.01036 | $k_1$ = -0.02417 |
| | $b_1$ = 0.295 | $b_1$ = 0.295 | $b_1$ = 0.4883 | $b_1$ = 0.2121 | $b_1$ = 0.4883 | $b_1$ = 0.295 | $b_1$ = 0.4883 | $b_1$ = 0.2121 | $b_1$ = 0.4883 |
| | $k_2$ = 0.029 | $k_2$ = 0.029 | $k_2$ = 0.01611 | $k_2$ = 0.01611 | $k_2$ =0.029 | $k_2$ = 0.145 | $k_2$ = 0.145 | $k_2$ = 0.145 | $k_2$ = 0.01611 |
| | $b_2$ = -0.865 | $b_2$ = -0.865 | $b_2$ = -0.4783 | $b_2$ = -0.4783 | $b_2$ = -0.865 | $b_2$ = -4.345 | $b_2$ = -4.345 | $b_2$ = -4.345 | $b_2$ = -0.4783 |
| Predation | $a_1$ = 1.464, $h_1$ = 0.01613; $a_2$ = 1.177, $h_2$ = 0.008982; $a_3$ = 1.437, $h_3$ = 0.01155; $a_4$ = 1.219, $h_4$ = 0.003985; $a_f$ = $a_m$ =1.461, $h_f$ = $h_m$ = 0.004453 | | | | | | | | |
| Fecundity | $Q_p$ = 100 | | | | | | | | |
